# Physiological responses of submerged freshwater macrophytes to multiple stressors

**DOI:** 10.64898/2026.03.23.713585

**Authors:** Amine M. Mahdjoub, Severin Eispanier, Elisabeth M. Gross, Sabine Hilt

## Abstract

- Submerged macrophytes are central to freshwater ecosystems functioning but are declining globally under multiple anthropogenic stressors. We aimed to identify general patterns in physiological responses and interaction types, and to assess whether a mechanistic understanding of stressor interactions can be developed from published evidence.
- We systematically reviewed 12,858 records, identified 172 relevant papers, and extracted effect sizes from 124 experiments included in the meta-analysis.
- Most studies examined combinations of nutrient enrichment, shading, toxic trace metals, warming, and emerging contaminants such as PFAS and microplastics, typically under simplified 2 x 2 factorial laboratory designs. Additive effects dominated (50%), while synergistic interactions were relatively infrequent (14%). Antagonistic interactions often reflected dominance of a single stressor or compensatory responses, whereas synergisms were most frequent with metals combined with co-stressors enhancing bioavailability.
- Our synthesis suggests that accumulated stressors cause negative, but not necessarily amplified, responses, although the limited number of experiments testing more than two stressors means synergistic effects may be underestimated. We propose *Stuckenia pectinata* as a model organism because of its cosmopolitan distribution, experimental tractability, and available genomic resources, and argue that expanding stressor complexity, duration, and taxonomic breadth will strengthen predictions of macrophyte responses and inform freshwater conservation under global change.

## 1. Introduction

Submerged macrophytes are key primary producers in freshwaters where they regulate ecosystem functioning through their influence on oxygen dynamics, nutrient cycling, habitat structure, and food web interactions (Jeppesen et al. 1998; Thomaz 2023). These ecosystem-level effects are ultimately governed by plant physiological performance, including the capacity to acquire carbon and nutrients, maintain photosynthetic efficiency, and allocate resources under environmental stress (Arnao et al., 2022; Gang et al., 2025). Despite their ecological importance, submerged macrophytes have undergone widespread and accelerating declines (Botrel & Maranger, 2023; Luo et al., 2025). Historically, these losses have been attributed primarily to eutrophication by anthropogenic nutrient loads (Botrel & Maranger, 2023; Phillips et al., 2016), in some cases, in combination with herbivory from water birds, herbivorous fish, crayfish and snails (Bakker et al., 2016; Hidding, 2016). However, recovery of submerged macrophytes has frequently failed even where nutrient inputs have been reduced (Hilt et al., 2006; Jeppesen et al., 2007), indicating that additional stressors constrain macrophyte performance beyond eutrophication and herbivory alone.

Under global change, submerged macrophytes in freshwater ecosystems are increasingly exposed to multiple, co-occurring stressors including increasing water temperatures (O’Reilly et al., 2003; Johnson et al., 2024), altered light regime associated e.g., with browning (Blanchet et al., 2022), chemical pollution such as pesticides (Schafer et al., 2012), and emerging contaminants including pharmaceuticals, per and polyfluoroalkyl substances (PFAS), microplastics and nanoparticles (Li et al. 2023). Each of these stressors can directly affect plant physiology by altering photosynthetic efficiency, respiration, oxidative balance, or nutrient stoichiometry. When stressors act simultaneously, their combined effects may be non-additive, producing synergistic or antagonistic responses that cannot be predicted from single-stressor experiments and may lead to unexpected ecological surprises (Lindenmayer et al., 2010; Myers, 1995). Such non-additive effects are expected to arise from physiological trade-offs, for example when investment in stress defence limits carbon assimilation and allocation for growth, or when stressor disrupt the same metabolic pathways, resulting in interacting responses (Zandalinas & Mittler, 2022). Physiological responses integrate environmental conditions over short timescales and provide mechanistic insight into stress effects on plants (Lamers et al., 2020). Traits related to photosystem performance, oxidative stress regulation, hormonal signalling, and resource allocation often respond more rapidly and sensitively than morphological or population-level endpoints (Arnao et al., 2022). Consequently, physiological measurements are particularly well-suited to detecting early stress responses and revealing the mechanisms underlying macrophyte decline under multiple stressors.

Although multiple-stressor research has expanded rapidly in freshwater ecology, submerged macrophytes remain largely overlooked. Most experimental and synthetic studies have focused on animals, especially fish and invertebrates, whereas macrophytes are rarely included (Bao et al., 2024; Birk et al., 2020; Jackson et al., 2016; Lange et al., 2018; Morris et al., 2022; Orr et al., 2024), A recent systematic review found that fewer than 1% of experimental freshwater multiple-stressor studies focused on submerged macrophytes, and even fewer examined physiological responses (Orr et al., 2024). Consequently, the nature and prevalence of their physiological responses to multiple stressors remain poorly understood. In contrast, studies on terrestrial and marine plants have generated substantial mechanistic insights into plant stress responses, often derived from experiments using well-established model species with fully sequenced and assembled genomes, such as the terrestrial model *Arabidopsis thaliana* (The Arabidopsis Genome Initiative 2000; Provart et al. 2016), or the marine *Zostera marina* (Davey et al., 2016; Stockbridge et al., 2020). These assembled genomes also allow the use of omics and the investigation of responses to stress at the molecular level (Zandalinas et al., 2021). However, it remains unclear whether such insights can be directly transferred to freshwater macrophytes, whether they share conserved stress-response pathways, or instead exhibit distinct, system-specific mechanisms. This knowledge gap is largely explained by the historical absence of high-quality genome assemblies for freshwater submerged macrophytes. This bottleneck is currently being overcome through initiatives such as The Darwin Tree of Life Project Consortium (2022) allowing assemblies also for these plants.

Here, we synthesise experimental evidence on the physiological responses of submerged freshwater macrophytes to multiple stressors. Using a meta-analytic approach, we compiled studies reporting physiological endpoints, and quantified responses to single and combined stressors. We classified stressor interactions as additive, synergistic, reversal, or antagonistic and quantified their relative frequency across stressor combinations. We predicted that research attention would be focused toward globally pervasive stressors such as warming, eutrophication, and pollution, and that although many combinations would yield additive effects, non-additive responses would arise when stressors impose convergent or conflicting demands on core physiological processes or when one stressor alters the physicochemical context governing the intensity, availability, or toxicity of another (Rider et al., 2014; Rillig et al., 2021). We further identify a suitable submerged macrophyte model species based on its distribution, functional traits, and genomic resource availability for stress gene family analyses. By disentangling dominant stressors and recurrent physiological trade-offs, this synthesis establishes a comparative framework for evaluating submerged macrophyte vulnerability under global change and defines priorities for mechanistically grounded experimental research.

## 2. Material and methods

### 2.1. Screening process and data collection

On February 13^th^, 2025, a literature search was conducted in the Web of Science (WoS) database using an advanced search query considering our four inclusion criteria of experiments studying: a fully submerged freshwater macrophyte species, at least a combination of two stressors tested, with at least one being experimentally manipulated, and their effects tested separately and in combination in comparison to a control treatment. A second search was conducted on December 3^rd^, 2025 to integrate studies published in 2025.

In this study, we used an adapted version of the stressor definition by Crain (2008), Piggott (2015) and Côté et al. (2016): A pressure resulting from anthropogenic activities, which induce an environmental change resulting in effects on organisms at the individual, population, or community level that may be either negative (disruption of homeostasis) or positive.

The resulting records were sorted using a machine learning classifier software (ASReview v2.0a11) to improve the process of finding relevant studies. To ensure the good quality and exhaustivity of the search, we followed the SAFE procedure (Boetje & van de Schoot, 2024), an active learning-based screening process. In Supplementary text 1, we describe the inclusion and exclusion process and the SAFE screening procedure, and in Fig. S1 we summarize these steps in a “Preferred Reporting Items for Systematic Reviews and Meta-Analyses” (PRISMA) like flowchart.

From each relevant record, experimental metadata were extracted using an adapted version of the stressor taxonomy developed by Orr et al. (2024), which was originally based on the terrestrial framework of Rillig et al. (2021) and subsequently refined for freshwater systems. We further modified this taxonomy to suit freshwater macrophyte systems. These data were analysed on R 4.4.3 to identify general trends and frequencies of studied stressors, model species, endpoint use in the literature.

### 2.2. Meta-analysis

From relevant studies that reported mean, variance and sample size, we extracted values from control treatment, single stressor treatments and combined stressors treatments, of macrophyte physiological endpoints that were measured in at least 10 experiments from different studies. We extracted values from tables, datasheets available in published supplementary material or directly from figures using WebPlotDigitizer (Version 5.2 Rohatgi 2024). If the standard error was reported, we calculated the standard deviation using: *SD* = *SE* × √*n* where *SD*, *SE* and *n* are the standard deviation, standard error and sample size, respectively. When values were available at multiple time points, we extracted data at the time point closest to 14 days, as most studies ran their experiments for 2 weeks.

#### 2.2.1. Effect size calculation

We calculated the natural log-transformed response ratio (lnRR) as an effect size measurement as it quantifies the proportion of change in stress treatments in comparison to the control treatment and has greater statistical power than other measures of effect sizes such as the standard deviation mean (Yang, Sánchez-Tójar, et al., 2023). We accounted for small sample size bias by applying corrections recommended in Lajeunesse (2015). We used equation (1) to calculate the effect size and equation (2) to calculate the sampling variance:

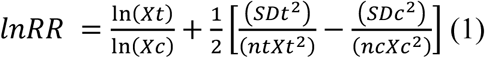

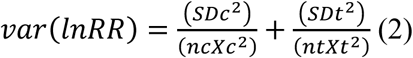

Where *X, SD* and *n* are the mean values, standard deviations and sample sizes of the measured endpoints of the stressor treatment *t* and control *c*. For an intuitive description and discussion of selected results, we calculated a percentage change (%) following equation (3):

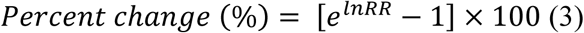

#### 2.2.2. Model fitting

We fitted a set of multilevel mixed-effects meta-analytic models to quantify how different physiological endpoints respond to single stressors, and compare how different stressor classes affect plant physiology. For single stress responses at the physiological endpoint level, we considered as random effects: the experiment, species, and stressor identity nested within each experiment, as stressor identities and applied levels (e.g., concentration or intensity) were tied to the experiment: *Effect size ∼ 1 +(1|Experiment) + (1|Species) + (1|Experiment/Stressor)*. For single stress response across all major endpoints, we first reversed the effect sizes sign of endpoints that are biologically known to increase with stress, mainly oxidative stress markers, so that negative values indicate reduced macrophyte performance and for the models we added physiological endpoint as an additional random effect: *Effect size ∼ 1 +(1|Experiment) + (1|Species) + (1|Physiological endpoint) + (1|Experiment/Stressor)*.

As experiments reported multiple single-stressor treatments sharing the same control, we constructed a variance-covariance (VCV) matrix per ‘study x species x control’ to meta-analyse endpoint changes with stress, and per ‘study x species x endpoint x control’ for modelled stressor estimates, following Lajeunesse (2011). We used the *rma.mv* function of *metafor* 4.8 package to calculate estimates and 95% confidence interval and considered estimates as significant if the confidence interval excluded 0. We quantified heterogeneity using *I²* (Higgins & Thompson, 2002), computed from the estimated variance components (*σ²*) and the mean sampling variance:

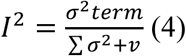

Where *I²* is the heterogeneity index ranging from 0 to 1, *σ*^2^ the term variances and *v* the within-study variance. To obtain the simplest efficient model, for each estimate, we began with all terms and dropped those with near-zero explained variance and thus heterogeneity (*I²* < 10^-6^), reflecting no information gain. The retained structure for each physiological endpoint and stressor class are reported in Tables S4 and S5. To test for publication and small study bias, we plotted funnel plots and performed Egger’s test via weighted least squares. Funnels were produced for all effect sizes including single and multiple stress treatments, and we report Egger p-values in Fig. S2.

#### 2.2.3. Nature of stressor interaction

We assessed the nature of interactions between pairs of stressors (2×2 designs) using a multiplicative null model. Under this framework, the expected effect of the combined treatment corresponds to the sum of effect sizes (lnRR) from single-stressor treatments, which represents an additive model in lnRR space and a multiplicative model on the original response scale. The observed combined effect was compared against this expectation. We classified interactions into four major categories following Crain et al. (2008): additivity, antagonism, synergism, and reversal. More precise interaction categories have been proposed by Piggott et al. (2015), however, their interpretation is less straightforward in applied and management-oriented contexts (Côté et al., 2016). However, the classification by Piggott et al. addresses important cases where responses to single stressor treatment act in opposing directions. Therefore, we applied a hybrid classification integrating both frameworks: if the difference between the expected and observed effect was non-significant, the interaction was classified as additive (Fig. 1a-c). We classified the interaction as synergistic if the observed effect was significantly more extreme than the expected effect (in absolute terms to account for effect sizes sign direction) (Fig. 1g-i) and antagonistic if it was significantly lower than expected (Fig. 1d-f). When single-stressor effects were of opposing sign, interactions were classified as synergistic if the combined effect exceeded the magnitude of the strongest single-stressor response, indicating that stressors did not cancel each other. Finally, when both single-stressor effects were of the same sign while the combined effect was significant in the opposite direction, the interaction was classified as a reversal.

**Fig. 1:**
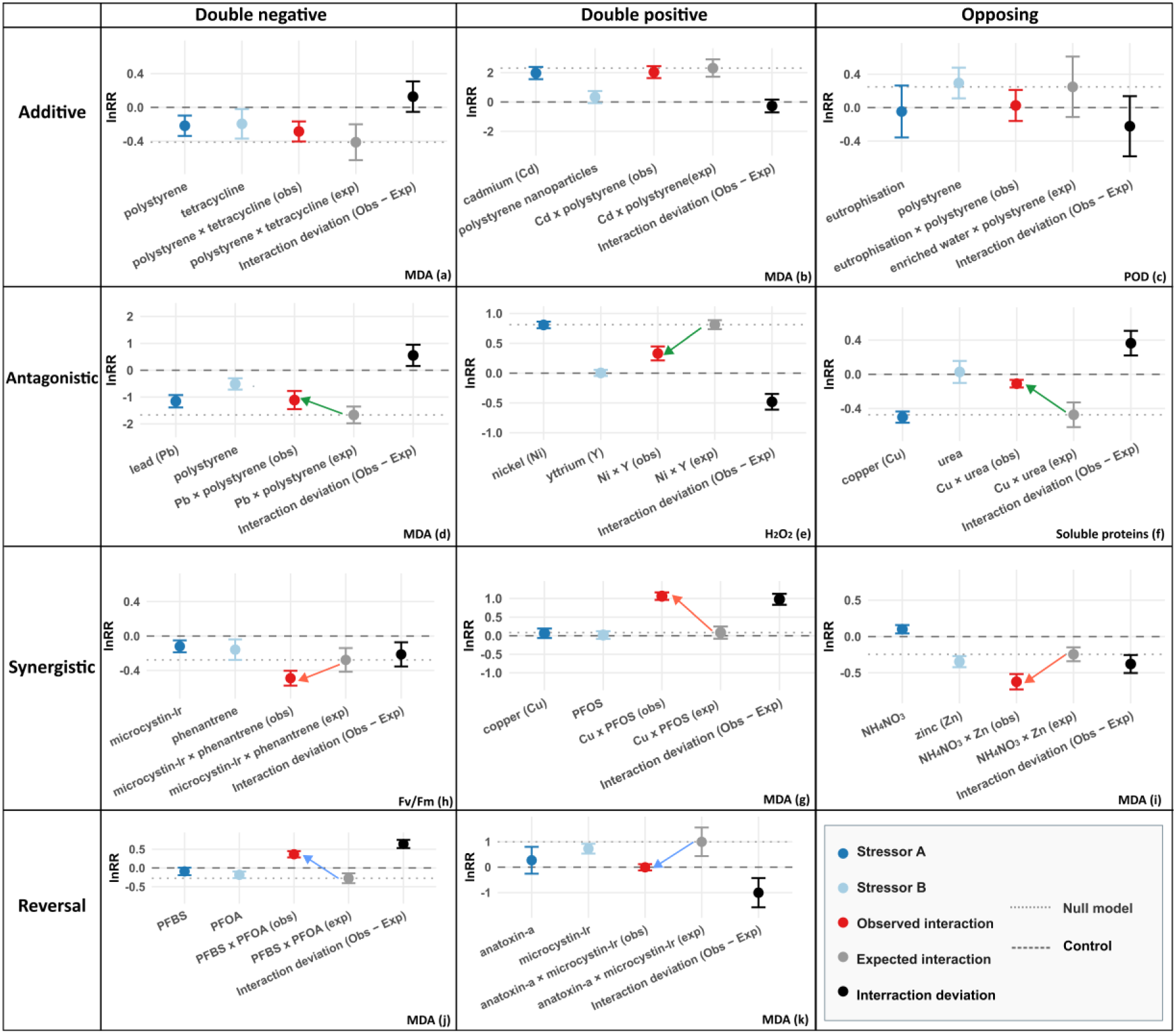
Examples of stressor interaction effects on different physiological endpoints in freshwater macrophytes. Endpoints shown include MDA (malondialdehyde), POD (peroxidase), CAT (catalase), and H_₂_O_₂_ (hydrogen peroxide), Soluble proteins, F_v_/F_m_ (maximum quantum yield). Panels correspond to the following studies: (a) Mao et al. (2024); (b) Wei et al. (2025); (c) Yu et al. (2022); (d) Ogo et al. (2022); (e) Lyu et al. (2019); (f) Maleva et al. (2016); (g) Wan et al. (2025); (h) Wang et al. (2023); (i) Wang et al. (2015); (j) Feng et al. (2025); (k) Li et al. (2024).

**Fig. 2:**
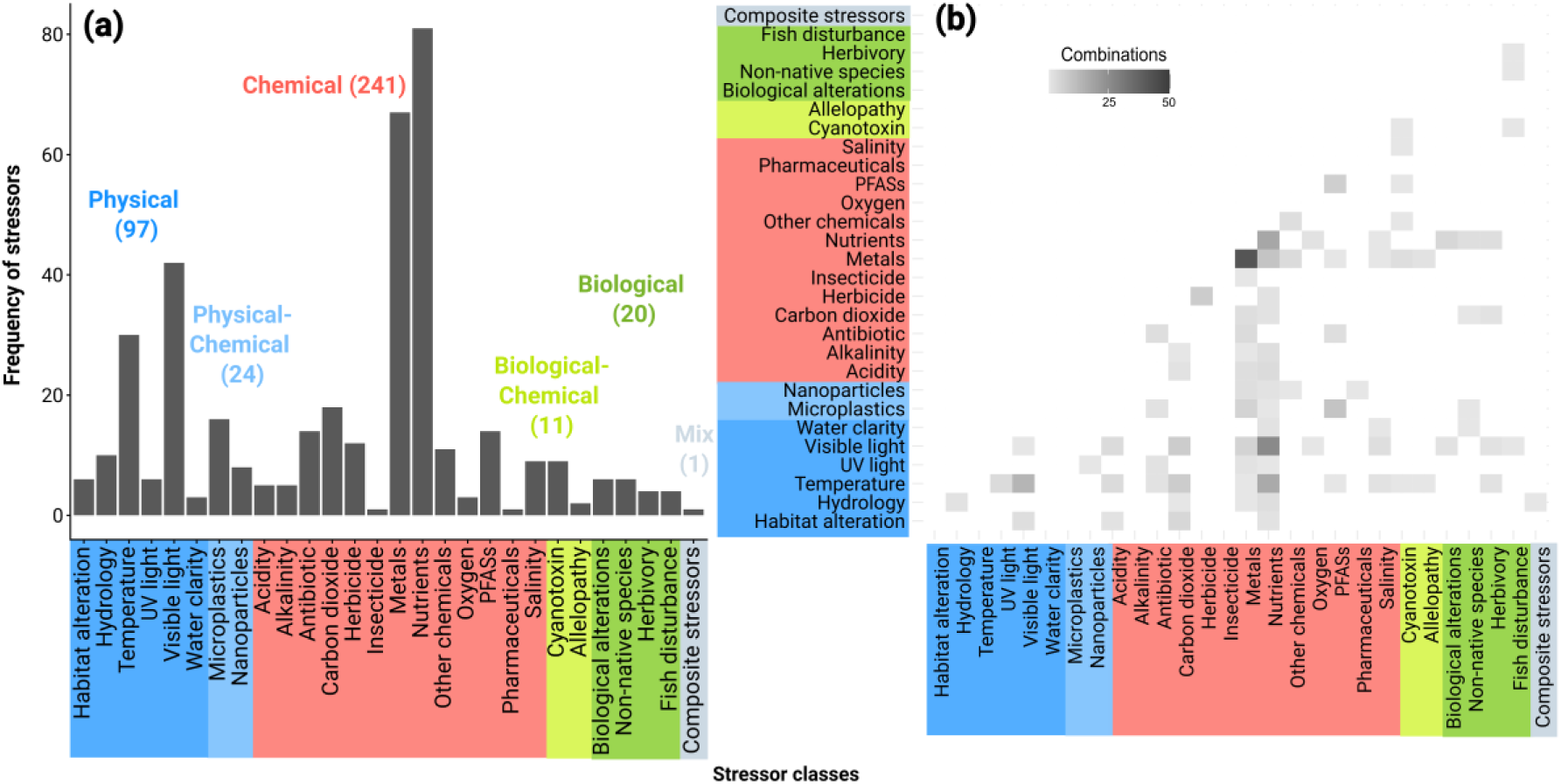
(a) Number of times a stressor class appears in a single stress treatment. (b) Frequency of interaction matrix for all multiple stressor treatments. Colours indicate stressor nature.

#### 2.2.4. Proteome prediction and phylogenetic analysis

After selecting *Stuckenia pectinata* as a potential model species (for details see below), we performed a comparative *in silico* investigation of copy number variation in oxidative stress related genes families using its new reference genome (GCA_976940525.1, The Darwin Tree of Life Project Consortium 2022). First, we generated an *ab initio* genome annotation using Tiberius v.1.1.8. We then used a comparative genomics pipeline to identify high-confidence candidate genes across five phylogenetically diverse plant relatives to facilitate phylogenetic footprinting, including: *Arabidopsis thaliana* as a reference, *Oryza sativa* and *Brachypodium distachyon* as monocot representative, *Zostera marina* as a marine relative, and *Wolffia australiana* as a floating freshwater species. We identified putative members of alternative oxidase (AOX) and Respiratory Burst Oxidase Homolog (RBOH) gene families by scanning for conserved protein domains (PFAM, PF01786, PF08414, PF08030, PF00175) using HMMER v.3.4 and filtered using stringent criteria (E-value <1e-10, bit score >50, length >100aa). High-confidence candidate genes were then independently validated by mapping to orthogroups containing canonical *A. thaliana* reference genes (Orthofinder v.3.1.2,). We compared gene counts across the selected species accessions (*A. thaliana* Araport11; *O. sativa* GCF_034140825.1; *Z. marina* GCA_001185155.1; *B. distachyon* GCF_000005505.3; *W. australiana* GCA_029677425.1). Detailed scripts, documentation and code used in this study are available at github.com/PHYTOPatCAU/Stuckenia_Stress.

### 2.3. Software use

We used R 4.4.3 and RStudio 2025.09.0+387 for data analysis and figure generation; details and references for all packages used are provided in Table S6. OpenAI ChatGPT 5 was used for limited refinement and troubleshooting of R code. Inkscape 1.4.2 was used for figure preparation and improvement.

## 3. Results

### 3.1. Trends and experimental designs

Between 1981 and 2025, 171 studies comprising 174 experiments investigated the physiological responses of freshwater macrophytes to multiple stressors, mainly conducted in China (54%). In parallel, 129 other studies focused solely on morphological trait measurements without including physiological data. Since the late 2010s, there has been a marked start of the use of molecular approaches, particularly metabarcoding of epiphytic biofilms, transcriptomics and metabolomics investigating gene expression, molecular pathways changes or alterations and metabolite production of aquatic plants (Fig. S2). A majority of experiments consisted of simple combinations with two stressor treatments (n = 139) and two levels tested, usually in the presence and absence of the stressor (Fig. S3a-c), in basically a 2×2 full factorial design. Gradient experiments with more than four stressor levels were rarely applied (n < 50) and the preferred experimental duration was between 10 and 30 days (n = 61), with 14 days most often applied (20 experiments). Shortest experiments were toxicity tests and ran for 1 to 3 days (n = 6) or as longer-term experiments of more than two months (n = 33), which were performed in outdoor mesocosm studies. Otherwise, indoor laboratory setting was prevalent (n = 112) and only two experiments were done in the field.

There were overall 127 unique stressors applied in 394 treatments across all experiments. Stressor treatments were largely of chemical nature (n = 226) followed by physical (n = 97) and only few of biological nature (n = 20) (Supplementary, Table S1). Four stressor classes were primarily tested and included nutrient enrichment (n = 81), toxic trace metals (n = 67), visible light reduction (n = 42) and temperature increase (n = 30) (Fig. 1a). The co-occurrence matrix (Fig. 1b) shows that popular stress interactions include metal toxicity cocktails (n = 42), combinations of nutrient enrichment and shading (n = 26), different types of nutrient manipulation such as water and sediment enrichment or different concentrations of phosphorus and nitrogen species (n = 18). Warming is also an emerging stressor often tested with nutrient enrichment (n = 17), or in combination with shading (n = 14). The matrix also reveals numerous gaps especially regarding biotic stressors.

Used species models were mainly angiosperms (30 out of 37), but also included four charophytes (*Chara vulgaris* L., *C. hispida* L., *C. braunii* S. G. Gmel. and *C. australis* R.Brown), two bryophytes (*Fontinalis antipyretica* Hedw. and *Jungermannia exsertifolia* Steph.) and one lycophyte (*Isoetes lacustris* L.). *Vallisneria natans* Lour. stands out in terms of frequency of usage, because experiments done in China usually selected it as a model (Fig. 3). We used the biogeographic realms classification of Udvardy (1975) to assess the representativity of different climatic and geographic regions (Fig. 3). The Palearctic realm, comprising Asia, Europe, and North Africa, was the most represented, whereas species occurring in the Afrotropical realm were comparatively scarce. Eight species were recorded across all realms, however, only four exhibit broad native distributions spanning multiple realms (*Ceratophyllum demersum* L., *Stuckenia pectinate* L., *Chara vulgaris* L. and *Potamogeton perfoliatus* L.), while occurrences in other regions likely reflect introductions beyond their native range.

**Fig. 3:**
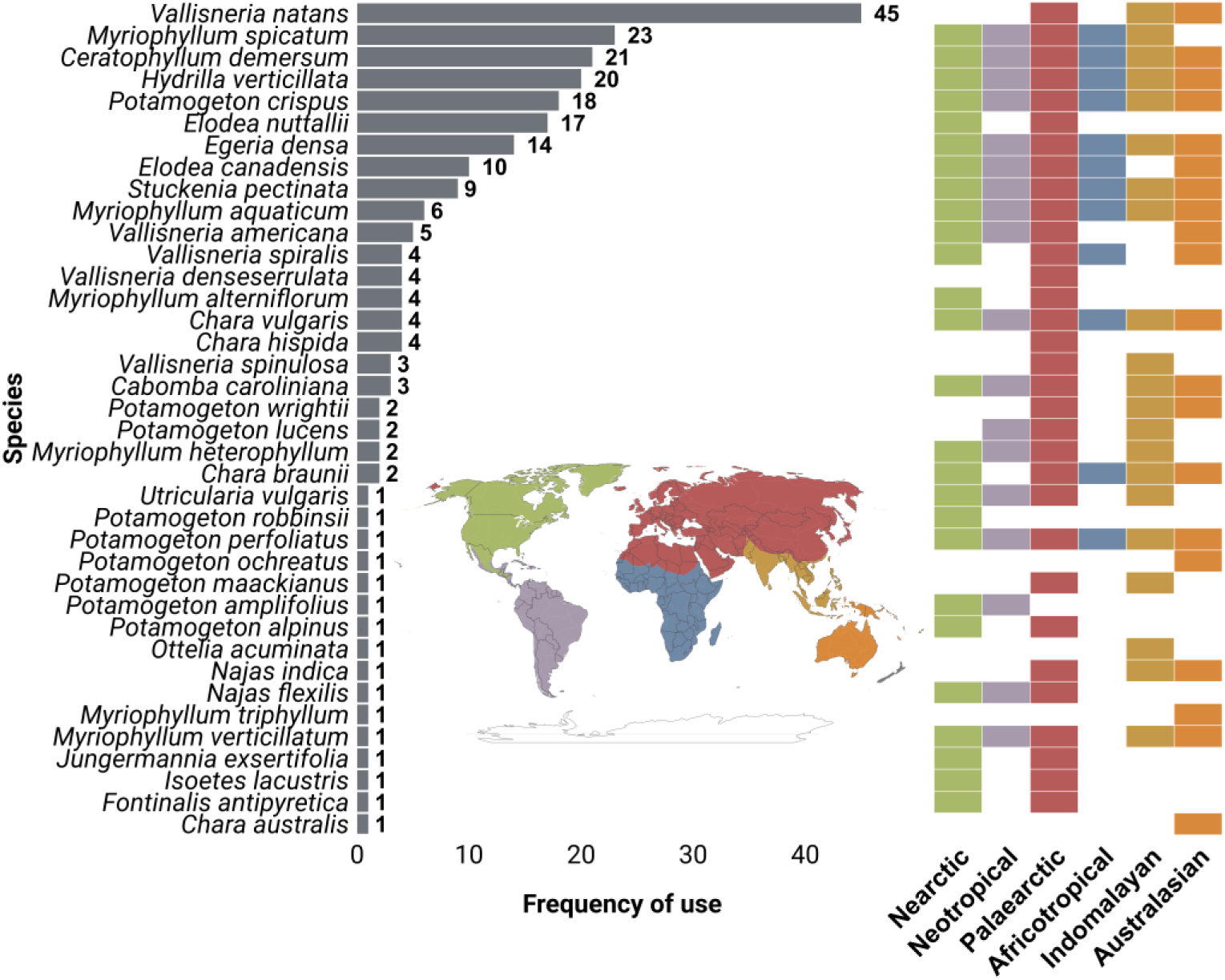
Frequency of used species and their biogeographic realm ranges according to the Udvardy (1975) classification.

Measured physiological endpoint classes primarily included photosynthesis-related traits, with pigments (chlorophyll a and b, carotenoids) often measured together, and chlorophyll fluorescence indices (the maximum quantum yield F_v_/F_m_ and the effective quantum yield Y) quantifying photosystem II photochemical efficiency, also commonly assessed together using a PAM (Pulse-Amplitude-Modulation) chlorophyll fluorometer (Fig. 4a). A second category included oxidative stress enzymes and markers, comprising malondialdehyde (MDA), catalase (CAT), peroxidase (POD), superoxide dismutase (SOD), and hydrogen peroxide (H₂O₂), typically assessed through biochemical assays. A third popular category concerned elemental stoichiometric status (carbon, nitrogen, and phosphorus). Regarding morphological measurements, most frequent traits included biomass and size changes as well as their relative variations through time expressed by relative growth rates (Fig. 4b).

**Fig. 4:**
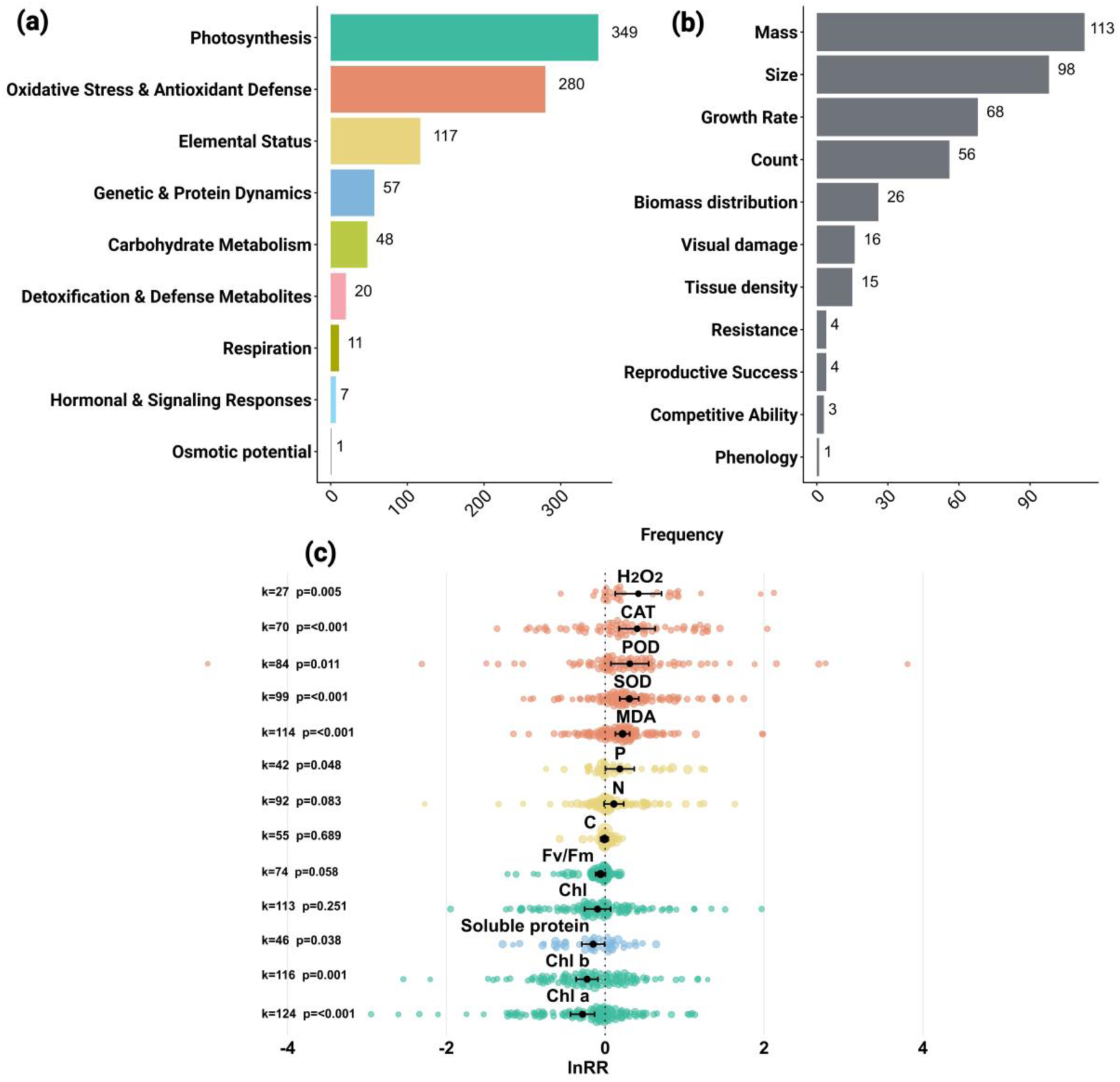
Frequency of measured physiological (a) and morphological (b) endpoint classes; detailed taxonomy is provided in Table S2. (c) Orchard plot showing physiological endpoint estimates (black points), 95% confidence intervals (black lines), and individual effect sizes (lnRR), with point size scaled by inverse standard error (1/SE). k indicates the number of effect sizes and p the p-value. Effects are not significant when confidence intervals cross the vertical dotted line.

### 3.2. Meta-analysis publication bias tests

The funnel plot of effect sizes (lnRR) of all treatments (single and multiple stress) shows a symmetrical distribution around zero, with most effect sizes clustering at low standard errors, indicating high precision (Fig. S5). Visual inspection suggests a weak asymmetry at higher standard errors where effect sizes are more dispersed towards negative values. Consistent with this pattern, Egger’s regression test indicated no evidence of publication bias (p = 0.062).

#### 3.2.1. Physiological response to single stress

Estimates of physiological changes modelled across all single-stressor treatments revealed distinct patterns tied to the nature of endpoints (Fig. 4c; Table S3). Among the 13 most frequently measured endpoints, 9 showed a significant response across all single stressor treatments. All oxidative stress markers (H₂O₂, CAT, SOD, POD and MDA) and phosphorus content increase with stress presence, while photosynthetic pigments (Chl a and Chl b), and soluble proteins decrease. Total chlorophyll given as Chl a + Chl b, nitrogen and carbon content, and the F_v_/F_m_ ratio do not change significantly.

The table in Fig. S4 details endpoint changes in response to each of the most often tested stressors, highlighting some class-specific stressor responses. While chlorophyll pigments decrease under most stressors, they increase with a reduction of light. Similarly, nitrogen content increases with light reduction and nutrient enrichment. Only two endpoints displayed consistent changes across stressors, and these were increased levels of oxidative stress markers and reduced F_v_/F_m_ ratios reflecting photosynthetic activity.

In the presence of a single stressor, the physiological performance of submerged macrophytes tends to decrease with the exception of shading and nutrient enrichment (Fig. 5). Toxic trace metals and PFAS had the most negative effects, while nanoparticles and microplastics also reduced plant performance. These were the only stressor classes showing a significant negative effect relative to controls, with consistent responses across endpoints (increased oxidative stress markers and decreased photosynthesis and elemental content). Other stressors had a marginal negative effect while an increase of carbon dioxide or nutrient enrichment barely had an estimated effect on macrophyte physiology.

**Fig. 5:**
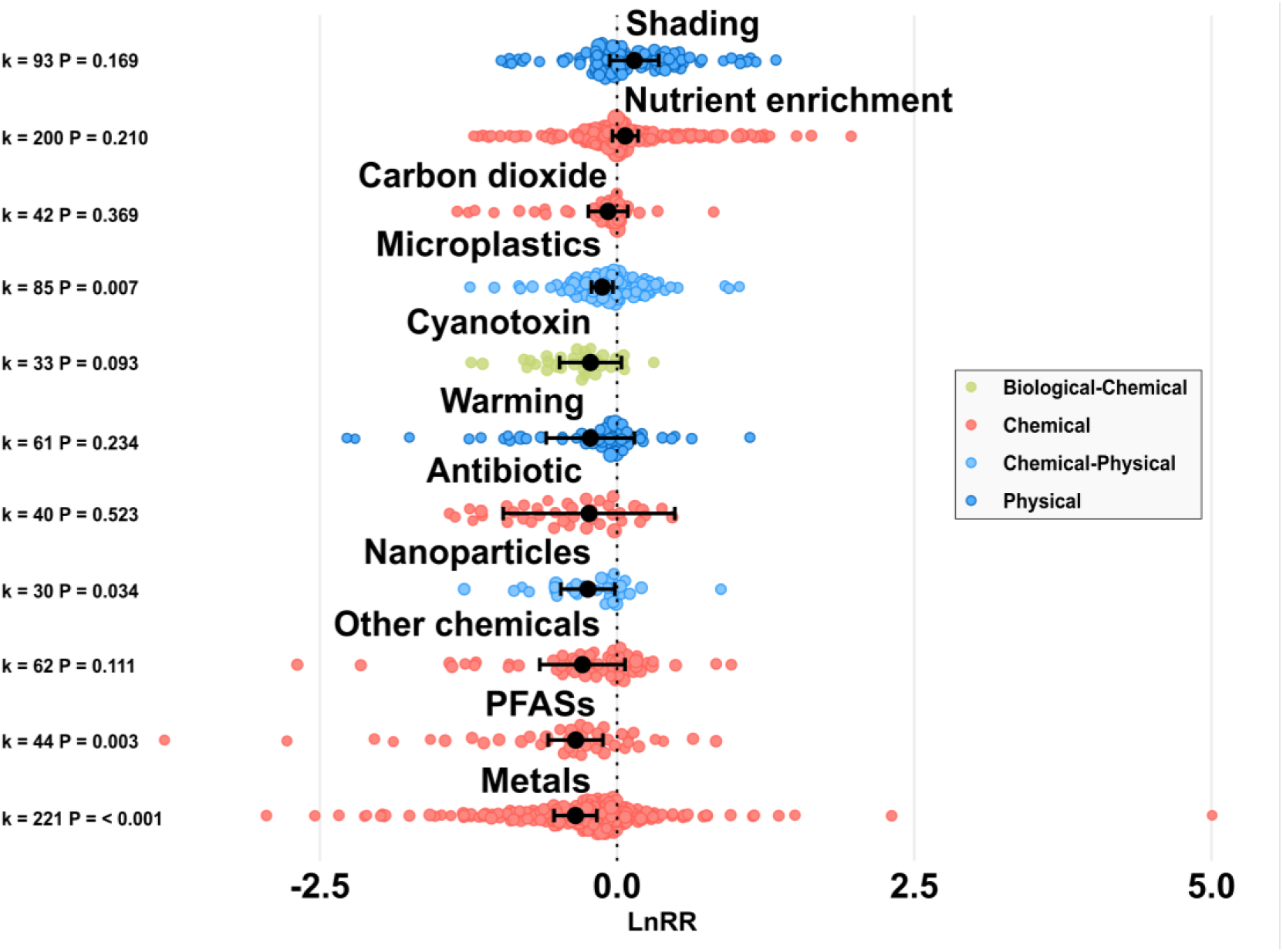
Orchard plot showing lnRR estimates (black points) of macrophyte physiological responses to single-stressor treatments by stressor class (10 most studied classes), with 95% confidence intervals (black lines) and individual lnRR effect sizes (points coloured by stressor nature).

#### 3.2.2. Physiological response to multiple stressors

A general assessment of physiological endpoint responses indicates that multiple stressors did not consistently amplify the direction or magnitude of responses compared with single-stressor effects (Fig. S5). Endpoints exhibiting negative responses under single-stressor exposure generally showed more negative estimates under multiple stressors, whereas endpoints responding positively to single stress tended to display more divergent response estimates when stressors were combined. Among the measured endpoints, none exhibited a statistically significant change under multiple stressors relative to single-stressor conditions (Fig. S5 - Welch’s unequal variances t-test – Table S5).

Simple combinations of stressors (2×2 factorial designs) induced overall 50% additive, 28% antagonistic, 13% synergistic, and 7% reversal interaction effects on submerged macrophyte physiology (Fig. 6). Interaction types were similarly distributed when single stressors induced effects in the same direction (i.e. double-negative or double-positive responses), with additivity being most frequent (44.6 and 44.3%), followed by antagonism (37 and 35.2 %), synergism (9.3 and 4.5 %), and reversal (8.9 and 15.9%). In contrast, additivity predominated in combinations (62.7%) involving opposing single-stressor responses with fewer ecological surprises (25.5% synergism and 11.7% antagonism). Despite the overall dominance of additivity, interaction class proportions were unevenly distributed across combinations and stressors, with additivity being almost ubiquitous (Fig. 6 and Fig. S6). Combinations involving metals had the highest proportion of synergism, whereas combinations involving PFAS mostly resulted in reversal and antagonistic interactions, particularly when mixtures included different PFAS species. Nutrient enrichment combinations were dominated by additive and synergistic interactions with no reversals. Combination including other chemicals or microplastics had a higher proportion of reversal, antagonistic, and synergistic interactions, leaving a relatively even distribution among these classes, while combinations including cyanotoxins were characterized by predominantly synergistic and reversal responses.

**Fig. 6:**
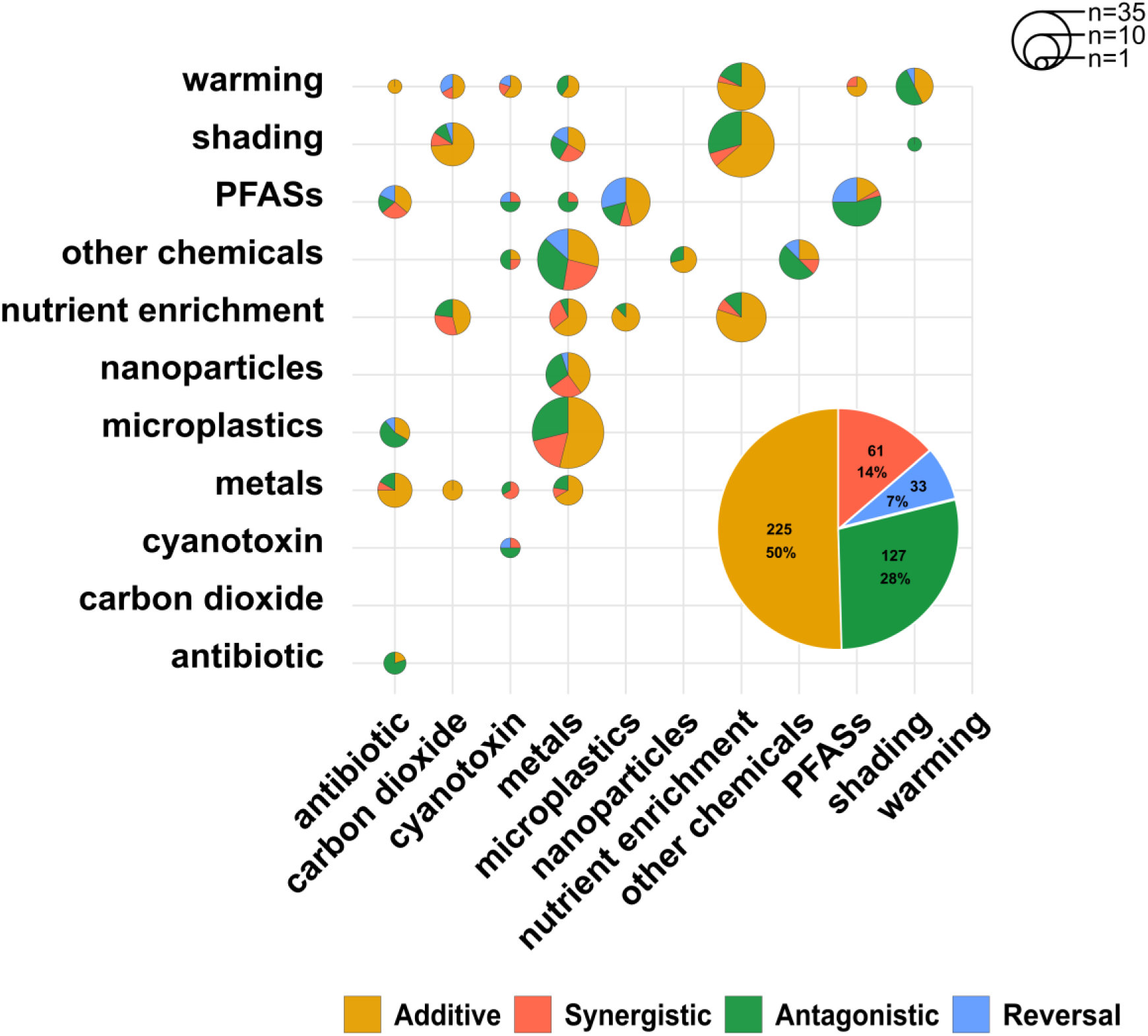
Scatter-pie chart showing the interaction types between the most frequently studied stressor pairs. Pie size reflects the number of effect sizes contributing to each pair, and pie segments indicate the proportion of additive, synergistic, antagonistic, and reversal interactions.

#### 3.2.3. Proteome prediction and phylogenetic analysis

Proteome predictions derived from the *S. pectinata* genome assembly enabled the identification and characterization of focal gene families involved in plant stress responses, allowing us to assess their representation in this submerged macrophyte. Comparative genomics analysis of the *S. pectinata* genome assembly revealed potential gene family expansions in oxidative-stress related gene families (AOX 7 genes vs. 5 in *A. thaliana* and 12 RBOH family genes vs. 10 in *A. thaliana*, Fig. S8). All identified gene duplications were supported by both stringent PFAM domain validation and orthogroup assignments, ensuring high confidence in our predictions. These findings are consistent with high BUSCO duplication rates on genome (D:37.0%) and the proteome-level (D:26.2%) reflecting the diploidized polyploid nature of *S. pectinata* (Fig. S9).

## 4. Discussion

This meta-analysis showed that simple stressor combinations in submerged freshwater macrophytes were mainly negative and additive, suggesting that stressor accumulation may accelerate population decline. Although some combinations showed antagonistic interactions, implying mitigating or unexpected protective effects, these patterns were often driven by dominance of one stressor, endpoint specificity, or combinations involving PFAS, microplastics, nanoparticles, or light reduction, all of which can modify co-occurring stressor effects. While theory predicts that synergistic interactions should account for approximately one third of interaction types (Côté et al., 2016), they represented only about one sixth in our dataset. However, the limited number of studies testing more than three stressors may underestimate their frequency and the potential for unexpected synergistic effects in natural systems (Zandalinas et al., 2021; Zandalinas & Mittler, 2022).

Our synthesis also revealed strong geographic, taxonomic, and experimental biases. Most studies were conducted in few regions, focused on few species, and used low replication with a limited number of stressors. Addressing these biases will require broader species and stressor coverage, higher replication, and experimental designs that better capture the complexity of natural freshwater ecosystems. This should improve prediction of how multiple stressors shape macrophyte physiology, population dynamics, and ecosystem functioning. In the following, we discuss the prevalence of tested stressor combinations and experimental designs, the occurrence of interaction types, the main measured endpoints, and the freshwater species most frequently studied.

### 4.1. Tested stressor combinations

Tested stressors in experimental studies largely reflect three major anthropogenic pressures on freshwater ecosystems. The first, and most frequently examined, is eutrophication, usually simulated through increased nutrient concentrations or manipulated light availability. Although this approach disentangles nutrient-enrichment effects linked to agriculture, more realistic designs should incorporate complex runoff scenarios, as in Vijayaraj et al. (2022), where cocktails of fungicides and pesticides were combined with nitrate enrichment, capturing full multiple stressors effect on freshwater ecosystems in agricultural catchments. The second major stressor group comprises chemical pollutants, historically dominated by trace metals but increasingly including microplastics, antibiotics, and PFAS. The third involves rising temperatures associated with global warming, which often interact with other stressors to alter macrophyte physiology. Most studies have focused on continuous warming, although recent evidence suggests that heat waves, rather than sustained warming, may more strongly exacerbate the effects of nutrient loading and herbicides on submerged macrophytes (Zhang et al. 2022).

Although these three stressor clusters also dominate experimental designs for other taxonomic groups (Orr et al., 2024), their prevalence and co-occurrence in the field remain poorly quantified. Other ecologically common stressors, including hydro-morphological alteration, sedimentation, and biotic pressures such as herbivory and pathogens, are underrepresented despite being frequent in lakes, streams, and rivers (Nõges et al., 2016). This is especially striking for biotic stressors, including plant pathogens, microbial community shifts, and invasive species, which are well documented in terrestrial plant systems (Agrios 2009; Suzuki et al. 2014a) but remain largely overlooked in freshwater plant research. As a result, the limited inclusion of these stressors constrains our ability to determine whether interactions observed in simplified experiments reflect natural systems or whether single stressors dominate macrophyte responses.

### 4.2. Experimental design

Most experimental studies on multiple stressors in submerged freshwater macrophytes rely on simplified 2×2 factorial designs, testing stressors at presence/absence levels with intensities selected based on environmental relevance (e.g. regulatory thresholds, field data, or predictive models). While these designs ensure feasibility and statistical power, isolating stressors under controlled laboratory conditions removes important ecological context. Key modulators, including hydro-morphology, water-column dynamics (particularly in lotic systems), natural temperature variability, and genotypic or ecotypic diversity, are rarely considered, and clonal identity is typically not controlled despite evidence that intraspecific variation can strongly influence stress tolerance and acclimation (Reusch, 2005; Roubeau Dumont et al., 2020). Moreover, such designs implicitly assume linear responses to stress intensity, although physiological responses are often non-linear and threshold-dependent, particularly for shading, warming and nutrients enrichment, where plants may exhibit optima or compensatory responses at moderate intensities but clear impairment beyond critical thresholds (e.g. >30 °C or >90% light reduction (Suzuki et al., 2014b).

Consequently, presence/absence designs may misclassify stressors as neutral or beneficial because responses depend on intensity. In addition, most experiments are short term, so transient physiological responses may obscure longer-term consequences such as altered population dynamics or increased vulnerability to subsequent stress. In natural systems, stressors can also interact indirectly through trophic or community-mediated pathways (Beauchesne, 2021). For example, pesticides may reduce periphyton grazers, promoting algal overgrowth and thereby increasing light limitation for macrophytes (Zhang et al. 2025). Such indirect and delayed interactions are largely absent from short-term factorial experiments, even though they may strongly shape outcomes in the field.

### 4.3. Multiple stressor interaction effect

Despite differences in study selection and interaction-classification frameworks, our results are consistent with previous meta-analyses of plant responses to multiple stressors. Additive effects also dominated in terrestrial plants and seagrasses (Rosenblatt & Schmitz, 2014; Stockbridge et al., 2020), accounting for 55% and 57% of cases, respectively. For seagrass, synergistic interactions were the next most common outcome (36%) and were driven primarily by combinations involving warming, whereas antagonistic interactions were rare (6%). Warming therefore deserves particular attention in freshwater systems, as it can modify the physical and chemical properties of co-occurring stressors and indirectly intensify pressures such as algal blooms, shading, nutrient competition, and herbivory, thereby promoting synergistic macrophyte decline.

In our dataset, antagonistic interactions often reflected stressor dominance or unchanged responses, regardless of whether individual stressors acted in the same or opposite directions. For example, shading-induced increases in pigment content could persist when combined with stressors that reduced pigments, producing antagonistic outcomes. Similar convergence on shared physiological pathways has been reported when single and combined stressors caused comparable reductions in antioxidant activity or photosynthetic performance (Chotikarn et al. 2021; Wang et al. 2021). Antagonism may also arise through interactions among stressor classes, as PFAS and microplastics can modify co-occurring stressors, alter uptake pathways, or provide substrate for microbial biofilms (Feng et al. 2024; Sun et al. 2025).

By contrast, combinations involving toxic trace metals, especially cadmium, arsenic, copper, and lead, most often produced synergistic interactions. Their strong individual effects, linked to ROS accumulation, impairment of antioxidant systems and chloroplasts, damage to Calvin-cycle proteins, and replacement of essential metals (Nagajyoti et al., 2010) may be further amplified by co-stressors that increase metal bioavailability or uptake. For instance, nanoparticles can adsorb metals and facilitate plant entry (Wei et al., 2025), nutrient enrichment may enhance metal sorption via biofilm and periphyton growth (Krayem et al., 2016), and acidification increases metal availability (Campbel & Stokes, 1985).

### 4.4. Measured endpoints

Abiotic and biotic stress challenge plant homeostasis, requiring coordinated phytohormonal and redox regulation to balance growth and defence (Arnao et al., 2022). Our meta-analysis shows that in submerged macrophytes, photosynthetic pigments and elemental status respond in a stressor-specific manner, reflecting adaptive or compensatory traits. Variables such as chlorophyll content and C, N, and P status are therefore useful to assess acclimation strategies, including shifts in resource allocation, but they are less direct indicators of stress than markers such as ROS, antioxidant compounds, or chlorophyll fluorescence. Focusing only on bulk C, N, and P changes, as is common in aquatic plant studies, also overlooks shifts in primary and secondary metabolism and within-plant resource reallocation.

Studies combining physiological and molecular approaches remain scarce (Fig. S2b) and are still largely descriptive compared with terrestrial systems, where cellular and molecular stress responses have long been investigated (Ebner, 2021; Rizhsky et al., 2002). Limited genomic resources have constrained most transcriptomic and metabolomic studies to untargeted treatment-versus-control comparisons. Nevertheless, transcriptomics has revealed broad differential expression under combined exposure to metals and PFAS, nanoplastics, or UV radiation, especially in pathways related to photosynthesis, antioxidant defence, carbohydrate metabolism, and signalling (Regier et al. 2016; Yi et al. 2020; Wang et al. 2023; Tang et al. 2024). Likewise, metabolomics has shown reprogramming of lipid, carbohydrate, and amino acid metabolism under PFAS, antibiotic, or cadmium exposure (Zhang et al. 2023; Yang et al. 2023; Y. Feng et al. 2024; S. Wang et al. 2024).

A smaller but growing body of studies has used metabarcoding to characterize submerged macrophyte biofilms, providing a first step toward describing the plant holobiont or meta-organism (Saha et al., 2024), and helping clarify microbial contributions to plant stress responses (Lopes et al., 2026). These biofilms are generally dominated by Pseudomonadota, and sometimes by Bacteroidota or Firmicutes, all common bacterial phyla on submerged plant surfaces (Wang et al. 2024). Responses to combined stressors appear context dependent, with several studies reporting declines in diversity under higher stress intensity or mixture treatments (Zhang et al. 2022; Hong et al. 2022; Yu et al. 2022; Li et al. 2023; Kang et al. 2024), whereas others report increases (Zhang et al. 2023; Yixia Yang et al. 2023; Huang et al. 2023).

Ultimately, molecular responses need to be linked to whole-plant performance to inform conservation, calling for integrative frameworks that combine transcriptomics, metabolomics, and microbiome analyses. Such approaches would enable macrophytes to be studied as meta-organisms, including the roles of endophytes in pollutant uptake or transformation (Weyens et al., 2009), nutrient cycling (Nair & Padmavathy, 2014), and the emergence of opportunistic pathogens under stress.

### 4.5. Tested freshwater plant species and proposal of a new model species

An ideal model species for freshwater macrophyte multiple-stressor research should be broadly distributed, easy to propagate under controlled conditions, show responses that generalize across systems, and ideally be supported by genomic resources. Against this background, the strong focus on *Vallisneria natans* remains limiting, as this species is restricted to Southeast Asia and is therefore not representative of global freshwater ecosystems. Future studies should broaden taxonomic and functional coverage by including globally distributed angiosperms such as the rooted *Stuckenia pectinata* and the free-floating *Ceratophyllum demersum*, as well as non-angiosperm groups such as bryophytes and charophytes, which play key ecological roles and are restoration targets (Blindow et al., 2014).

*Stuckenia pectinata* is particularly attractive as a model because it is experimentally manageable: it can be cloned by rhizome cuttings, propagated by tubers, and occurs across freshwater and brackish habitats, enabling comparisons with close relatives such as seagrasses (e.g. *Zostera marina*). Its ability to produce tubers is a key advantage, as these storage organs support establishment and regrowth under experimental conditions (Van Wijk, 1989). facilitating conditioning at experiment start. Moreover, *S. pectinata* tolerates shade and eutrophication (Kantrud, 1990) and can persist or even expand under conditions that cause other macrophytes to collapse, highlighting its conservation relevance (Hilt et al., 2013, 2018). Recent freshwater macrophyte genome assemblies, notably through the Darwin Tree of Life Project, now also include *S. pectinata* (The Darwin Tree of Life Project Consortium, 2022) enabling gene-expression and molecular studies that could further establish it as a model system. Its comparatively complex genome architecture, rather than being a drawback, may even be informative, potentially reflecting repeated transitions between land and water (Provart et al. 2016; Yang et al. 2020).

Building on these genomic resources, we used the *S. pectinata* assembly to identify candidate molecular targets involved in stress tolerance, with a focus on oxidative-stress pathways. Our comparative analyses suggested lineage-specific expansion of the AOX and RBOH gene families through duplication. Although some inflation during gene prediction cannot be fully excluded, stringent filtering and orthology-based curation support the biological plausibility of these patterns, further supported by BUSCO profiles indicating substantial duplication and proposed ancient polyploidy. Overall, these results provide a basis for future functional work and suggest that gene duplication may have contributed to oxidative-stress responses in *S. pectinata*.

## Conclusions

This study addresses how submerged freshwater macrophytes, key contributors to ecosystem stability and water quality, respond to single and multiple stressors. As freshwater systems face rising temperatures and increasing agricultural, industrial, urban, and biotic pressures, these plants are likely exposed to complex and sometimes unpredictable effects.

Our synthesis indicates that stressor accumulation mainly produces additive negative effects on macrophyte physiology and associated biofilm communities, although non-additive interactions remained common. However, the narrow range of stressor combinations tested limits inference under realistic conditions, where multiple stressors often co-occur and interact simultaneously (Nõges et al., 2016). In addition, many field-relevant stressors remain understudied, as most experimental work has focused on runoff-related pressures while many emerging threats are still poorly explored.

To improve mechanistic understanding and conservation relevance, future research should identify representative model species, and we suggest that the globally distributed and experimentally tractable *Stuckenia pectinata* could serve this role. Further progress will require higher replication, broader endpoint coverage, longer experiments, and more realistic multitrophic and hydrodynamic designs. This will strengthen understanding of cumulative stressor effects and support targeted management and restoration.

## Supporting information

Supplementary materila

## 5. Acknowledgements

This work was supported by the German Research Foundation (HI 1380/10-1) through the “PlantsCoChallenge” Research Unit 5640 (“Physiological and Evolutionary Adaptation of Plants to Co-occurring Abiotic and Biotic Challenges”). A special thanks to Thomas Mehner and Kate Laskowski for their valuable input during the courses on Scientific Writing and on General Linear Mixed Models.

## 7. Author contributions

**Conceptualization:** Sabine Hilt, Elisabeth M. Gross, Amine M. Mahdjoub. **Methodology:** all authors. **Investigation:** all authors. **Formal analysis and data curation:** Amine M. Mahdjoub, Severin Eispanier. **Visualization:** all authors. **Writing – original draft preparation:** all authors. **Writing – review and editing:** all authors. **Supervision, resources, project administration, and funding acquisition:** Sabine Hilt.

## 8. Data availability

The dataset of multiple-stressor studies is available in Dataset S1. The R scripts used to perform the meta-analysis are provided in the ZIP folder accompanying Dataset S2.

## 9. Supporting Information

The following Supporting Information is available for this article:

- **Supplementary text S1**. Screening process.
- **Fig. S1.** PRISMA-like flowchart of the literature screening and study selection process.
- **Fig. S2.** Trends in endpoint types and molecular approaches used in multiple-stressor studies on submerged macrophytes.
- **Fig. S3.** Temporal and geographic distribution of experiments included in the review.
- **Fig. S4**. Summary of experimental design characteristics across included studies.
- **Fig. S5.** Funnel plot of lnRR effect sizes and assessment of publication bias.
- **Table S1**. Stressor taxonomy used to classify anthropogenic pressures studied in multiple-stressor experiments on freshwater submerged macrophytes.
- **Table S2.** Endpoint taxonomy used to classify physiological and morphological measurements.
- **Table S3.** Summary of modelled physiological endpoint changes under single stress.
- **Table S4.** Summary of modelled physiological responses of submerged macrophytes to single stress.
- **Fig. S6.** Physiological responses of submerged macrophytes to the main single-stressor classes.
- **Fig. S7.** Comparison of lnRR estimates for frequently measured physiological endpoints under single and multiple stressors.
- **Fig. S8.** Distribution of interaction types across major stressor categories.
- **Fig. S9**. Comparative abundance of AOX and RBOH gene families across selected angiosperms.
- **Fig. S10.** Maximum-likelihood phylogeny of putative AOX and RBOH protein sequences.
- **Table S5.** R packages used for information synthesis, meta-analysis, and figure production.
- **References S1**. References for studies included in the review.

