## Supplementary materila for "Physiological responses of submerged freshwater macrophytes to multiple stressors"

**Supporting Information:** Physiological responses of submerged freshwater macrophytes to multiple stressors

**Methods S1. Screening process**

Two literature searches for experimental studies focused on the response of freshwater macrophytes to multiple stressors were done on *Web of Science.* A first one on 13 February 2025, and a second on December 03^rd^ 2025 to include papers published in 2025 using the following query:

*TS = (“macrophyte*” OR “submerged aquatic plant*”)
AND
TS = (“stressors” OR “factors” OR “drivers” OR “pulses” OR “synerg*” OR “antagon*” OR “additive” OR
“interact*” OR “inhibit*” OR “mediate” OR “combin*” OR “cumulative” OR “mixture” OR “factorial”)
AND
TS = (“freshwater*” OR “limnol*” OR “river*” OR “stream*” OR “lake*” OR “pond*” OR “wetland*” OR
“creek*” OR “aquatic”)
AND
TS = (“impact*” OR “effect*” OR “experiment” OR “mesocosm*” OR “microcosm*”)
AND
PY = (1900-2025)*

This query targeted studies that focused on freshwater submerged macrophytes, examined at least two anthropogenic stressors and their interactions, quantified biological responses to both individual and combined stressor effects, and were published between 1900 and 2025. The resulting records, including metadata (titles, abstracts, authors, and year of publication), were exported for further screening.

We used the machine learning tool ASReview (v2.0a11 for the Web of Science search conducted on February 13, 2025, and v2.2rc4 for the second search on December 3, 2025) to accelerate study selection. The software enables flexible model configurations, including the choice of classifier (algorithm), feature extractor (text analysers) and the specification of initial relevant and irrelevant records as prior data for model training. To ensure a rigorous and reproducible screening process, we followed the SAFE procedure (Boetje and van de Schoot 2024). This provides a standardized framework for semi-automated screening, defining a heuristic stopping criterion that ensures nearly all relevant studies are identified while minimizing unnecessary screening effort. Chosen stopping rule for all screening phases was that at least 20% of the records were screened, and screening was stopped after 50 consecutive records were labelled as irrelevant.

Phase 1: We first used a simple and fast configuration with a Naïve Bayes classifier and a TF-IDF feature extractor, based on the frequency of words in titles and abstracts. From the Web of Science query results and from the previously published multiple stressor database J. Orr et al. (2024), we selected 15 relevant and 15 irrelevant records to train the initial model.

Phase 2: We then switched to a configuration based on Logistic Regression (LR) combined with Sentence-BERT (SBERT), using the relevant and irrelevant records identified in Phase 1. This configuration has been reported to improve retrieval performance after a model switch (Teijema et al. 2022). Although it requires more prior data and computational power, it allows the model to identify additional relevant records more effectively.

Phase 3: We screened the records previously labelled as irrelevant to check for potential manual misclassification under the simpler Naïve Bayes + TF-IDF model configuration.

Phase 4: For all relevant records, we performed citation chasing using the CitationChaser tool (Haddaway et al. 2021), extracting both the references cited in the included studies and the studies citing them. This step thus also allows to identify studies not indexed in the Web of Science database. The resulting records were cleaned by removing books, reviews, non-English publications, and duplicates of previously identified or screened studies. The remaining records were then separated into those with abstracts and those without. Records with abstracts were screened using ASReview (Naïve Bayes + TF-IDF), whereas records without abstracts were screened manually based on title only.

Numerical details of the screening process is displayed in the following PRISMA style flow-diagram in Fig. S1.

**Identification of studies via other methods**

**Identification of studies via databases and registers**

Records identified from:

J. Orr et al., 2024 (n = 72)

Phase 4: Citation chasing analysis: (n = 8173)

Records identified from:

**Web of Science (ISI)**

Search on 13.02.2024 (n = 4206)

Search on 03.12.2025 (n = 479)

**Identification**

Records excluded

Manually excluded (n = 721)

Non-labelled (n = 3515)

Phase 1: active screening with Naïve Bayes-TF-IDF classifier (n = 4685)

Reports sought for retrieval

(n = 8246)

**Screening using ASReview and full text checks**

Phase 2: Deep learning: LR-SBert classifier (n = 3515)

Records excluded

Manually excluded (n =187)

Non-labelled (n = 3319)

Records with abstracts screening using 0ïve Bayes-TF-IDF classifier (n = 3490)

Records without abstract screened by title then full text skimming (n = 3992)

Reports excluded:

Non-English (n = 21)

Not articles (n = 303)

Duplicates (n = 54)

Manually excluded (n = 4255)

Non-labelled (n = 3543)

Phase 3: screening for mislabelled records (n = 908)

Records excluded

Manually excluded (n = 105)

Non-labelled (n = 828)

Physiological studies (n = 172)

Morphological only studies (n = 132)

**Included**

Fig. S1: PRISMA-like flowchart of our screening process. Original diagram source: Page MJ, et al. BMJ 2021;372:n71. doi: 10.1136/bmj.n71.

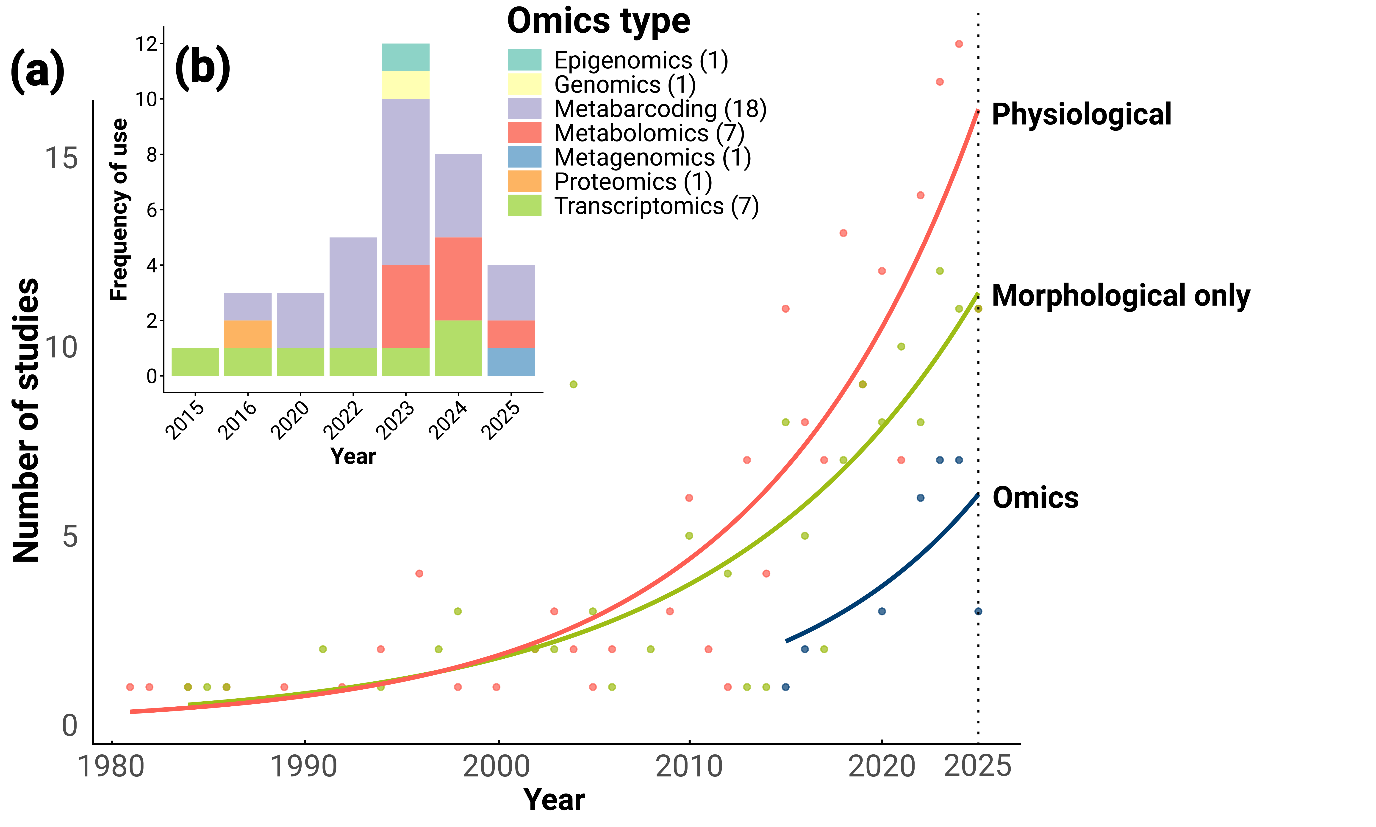

Fig. S2: (a) Trends of submerged macrophyte endpoint measurement nature in multiple stressor studies (physiological, morphological and omics). (b) Frequency of different types of molecular analysis performed.

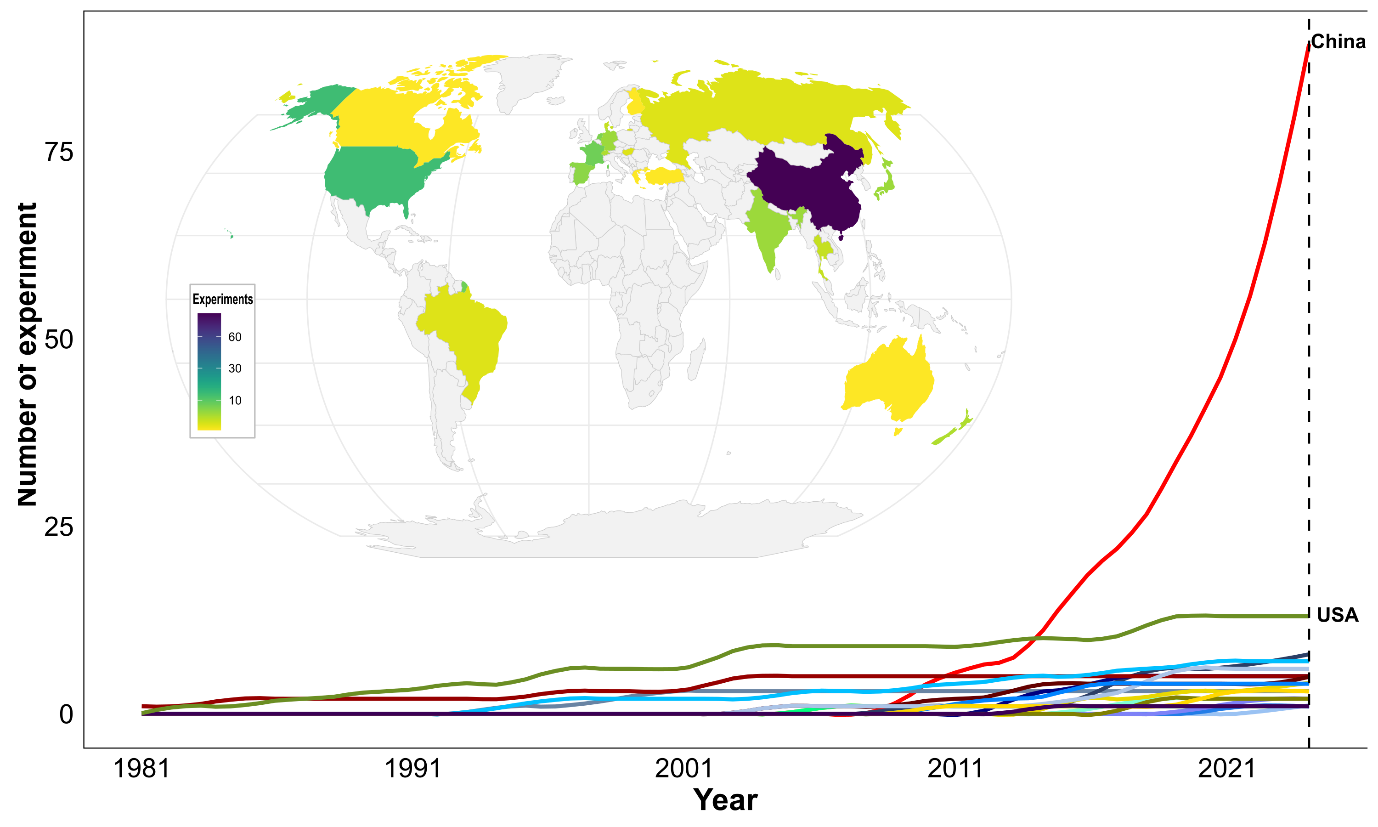

Fig. S3: (a) Cumulative frequency of experiments throughout the years, per country. (b) Global distribution of experiment location.

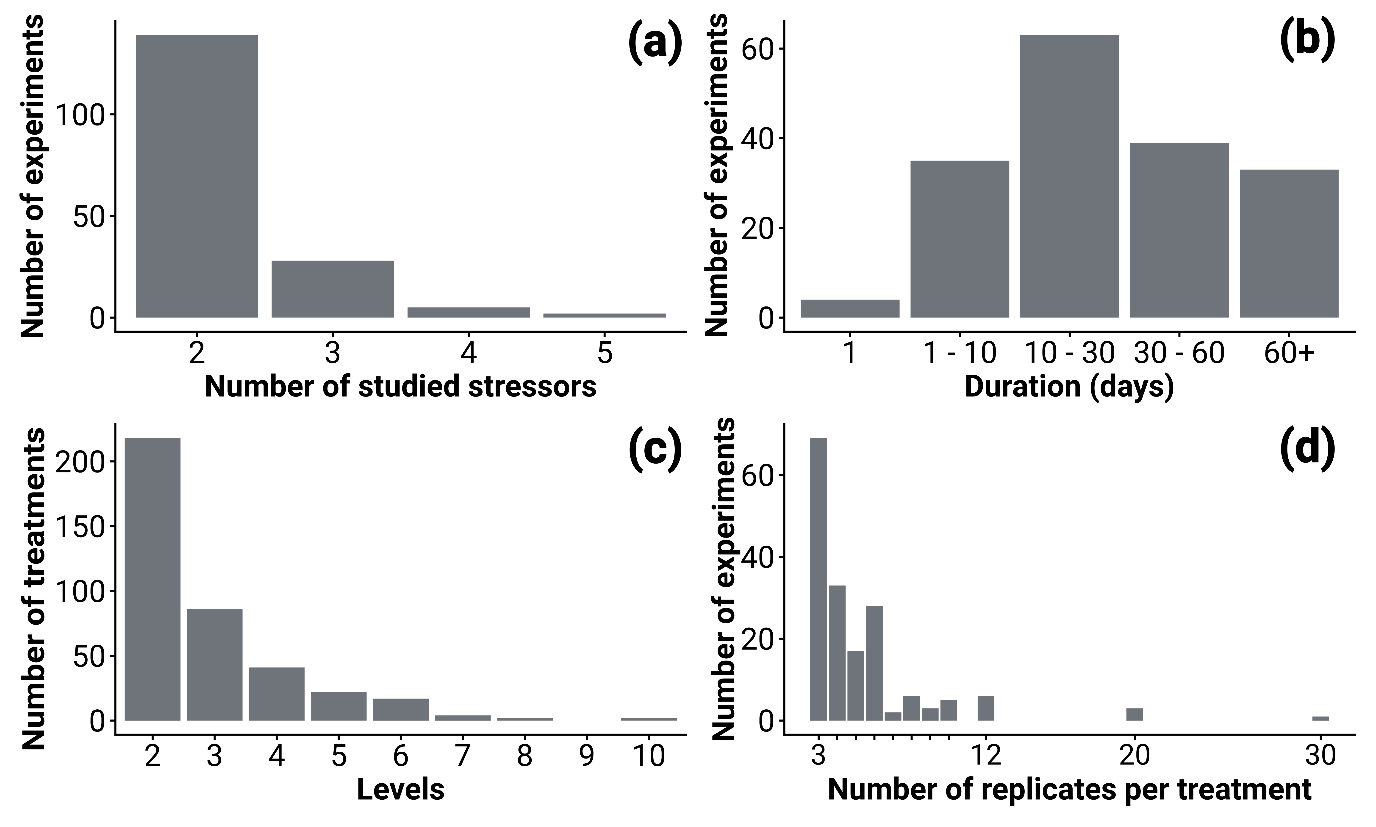

Fig. S4: Experimental design summary. (a) Complexity of experimental combinations. (b) Number of stressor treatments tested at different level (e.g.: presence/absence or high and low concentration). (c) Frequency of experiment duration. (d) number of experiments that used different number of used individual plants replicate.

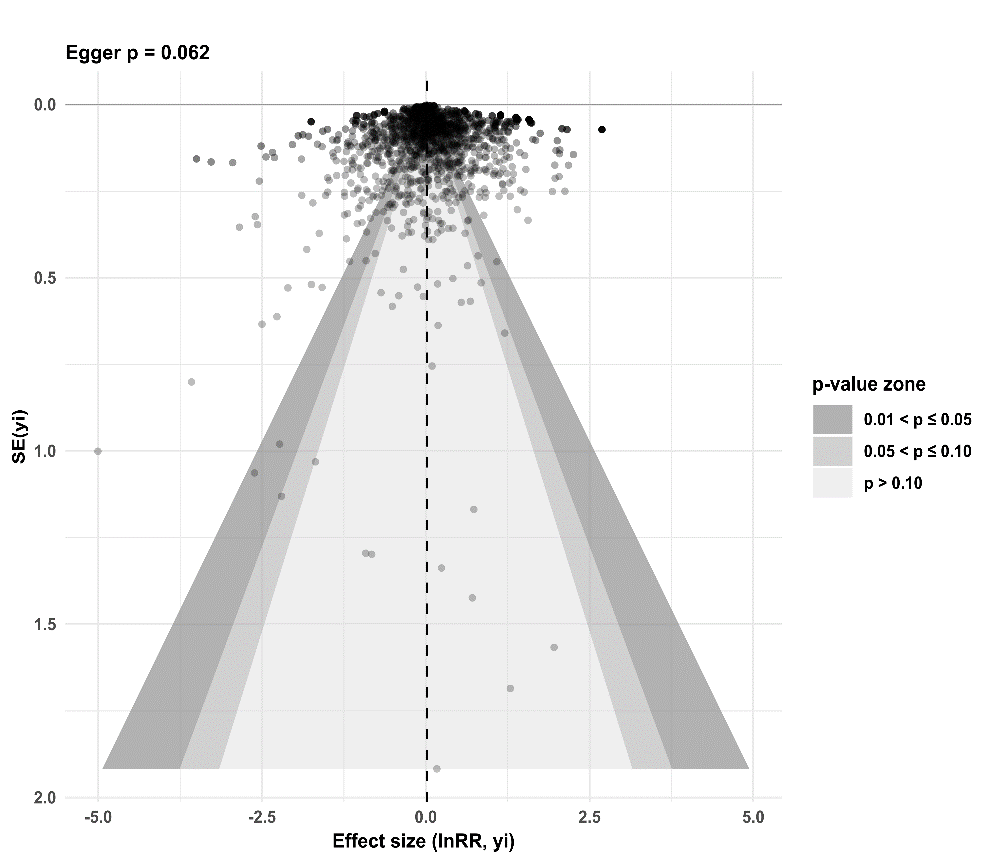

Fig. S5: Funnel plot of all effect sizes (lnRR) and assessment of publication bias with Egger’s test. SE: Standard Error.

Table S1: Stressor taxonomy used to classify anthropogenic pressures studied in multiple stressor experiments on freshwater submerged macrophytes.

| **Nature** | **Class** | **Identities** |
| --- | --- | --- |
| Physical | habitat alteration (6) | sediment type (5), sedimentation (1) |
|  | hydrology (10) | flow regime (1), flow velocity (4), water level (5) |
|  | temperature (30) | cooling (1), heat wave (1), temperature range (1), warming (27) |
|  | UV light (6) | UV light (5), UVB (1) |
|  | visible light (42) | dye (1), light colour (3), light intensity (5), photoperiod (1), shading (32) |
|  | water clarity (3) | brownification (1), depth (1), suspended particles (1) |
| Chemical-Physical | microplastics (16) | UVA-exposed polylactic acid (1), microplastic fragments (2), polyethylene (2), polyethylene terephthalate (1), polylactic acid (1), polypropylene (1), polystyrene (4), polyvinyl chloride (3), recycled polyvinyl chloride (1) |
|  | nanoparticles (8) | copper nanoparticles (2), gold nanoparticles (1), nanoplastics (3), polystyrene nanoparticles (1), titanium dioxide nanoparticles (1) |
| Chemical | acidity (5) | humic acid (1), low pH (3), pH range (1) |
|  | alkalinity (5) | bicarbonate (3), hardness (1), high pH (1) |
|  | antibiotic (14) | anrofloxacin (1), antibiotic mix (1), azithromycin (1), chloramphenicol (1), enrofoxacin (1), erythromycin (1), oxytetracycline (1), sulfadiazine (2), sulfamethoxazole (1), tetracycline (4) |
|  | carbon dioxide (18) | carbon dioxide range (4), increased carbon dioxide (14) |
|  | herbicide (12) | aminomethylphosphonic acid (1), asoproturon (2), atrazine (2), diuron (2), glyphosate (1), isoproturon (1), isothiazolinon (1), metsulfuron-methyl (1), sodium dichloroisocyanurate (1) |
|  | insecticide (1) | hexachlorocyclohexane (1) |
|  | metals (67) | arsenic (6), cadmium (20), chromium (2), copper (18), iron (2), lead (4), manganese (2), mercury (3), metal mixture (1), nickel (4), yttrium (1), zinc (4) |
|  | nutrients (81) | ammonia (3), ammonium (13), ammonium nitrate (2), dissolved organic matter (1), enriched sediment (12), enriched water (20), nitrate (8), nitrogen (6), nutrient enrichment (1), phosphate (2), phosphorus (9), potassium (3), sucrose (1) |
|  | other chemicals (11) | benzaldehyde-A (1), benzaldehyde-M (1), bisphenol f (1), chloroacetic acid (1), n-nitrosodimethylamine (1), phenanthrene (1), phenol (1), sodium dodecyl sulfate (1), sodium nitroprusside (1), urea (2) |
|  | oxygen (3) | low oxygen concentration (2), sediment anoxia (1) |
|  | PFASs (14) | PFASs mix (1), hexafluoropropylene oxide dimer acid (2), perfluorobutane sulfonate (1), perfluorobutyric acid (2), perfluorooctane sulfonate (1), perfluorooctanesulfonic acid (1), perfluorooctanoic acid (6) |
|  | pharmaceuticals (1) | diclofenac (1) |
|  | salinity (9) | increased salinity (8), sodium chloride (1) |
| Biological-Chemical | cyanotoxin (9) | *Microcystis aeruginosa* (3), anatoxin-a (1), microcystin-lr (5) |
|  | allelopathy (2) | *Eichhornia crassipess* allelopathic substances (1), exogenous anthocyanin extract (1) |
| Biological | biological alterations (6) | mowing (2), periphyton (4) |
|  | non-native species (6) | *Alternanthera philoxeroides*, *Myriophyllum aquaticum* (1), *Elodea nuttallii* (2), *Hydrilla verticillata* (1), *Myriophyllum alterniflorum* (1), non-native species (1) |
|  | herbivory (4) | *Bellamya aeruginosa* (1), *Radix swinhoei* (3) |
|  | fish disturbance (4) | *Misgurnus anguillicaudatus* (3), *Pseudorasbora parva* (1) |
| Mixed | composite stressors (1) | copper and nitrogen (1) |

Table S2: End point taxonomy used to classify physiological and morphological measurement used.

| **Nature** | **Class** | **Identity** |
| --- | --- | --- |
| Physiological | Carbohydrate Metabolism | invertase (1), pyrophosphatase (ppi) (1), soluble carbohydrates (10), soluble sugars (15), starch (17), sucrose (1), total carbohydrates (3) |
|  | Detoxification & Defence Metabolites | flavonoids (2), metallothionein (1), phenylalanine ammonia lyase (1), phytochelatins (1), polyphenol oxidase (1), proanthocyanidins (1), total phenolic compounds (10), ultraviolet radiation absorbing compounds (2), urease (1) |
|  | Elemental Status | calcium content (2), carbon content (30), magnesium content (3), nitrogen content (44), organic matter content (1), phosphorus content (28), potassium content (6), potassium leakage (1) |
|  | Genetic & Protein Dynamics | cysteine (4), dna content (1), free amino acid content (15), glutamine (1), protease (1), rna content (1), rnase (1), s-adenosylmethionine decarboxylase (adometdc) (1), total soluble protein content (31), glutamine synthetase activity (1) |
|  | Hormonal & Signalling Responses | abscisic acid (2), indole-3-acetic acid (iaa) (1), indoleacetic acidâ oxidase (1), jasmonic acid (1), nitric oxide (1), strigolactone (1) |
|  | Oxidative Stress & Antioxidant Defence | ascorbate peroxidase (8), ascorbic acid (4), catalase (CAT) (39), diamine oxidase (1), docosahexaenoic acid (DAO) (2), glutathione (16), glutathione s-transferase (GST) (9), glutathione disulfide (1), glutathione peroxidase (GPX) (3), glutathione reductase (5), glycolate oxidase activity (1), hydrogen peroxyde (H2O2) (15), malondialdehyde (MDA) (57), ornithine decarboxylase (1), peroxidase (POD) (43), proline (10), superoxide (O2^-^) (7), superoxide dismutase (SOD) (54), total reactive oxygen species (ROS) (1), total acid soluble thiols (3) |
|  | Photosynthesis | antheraxanthin (1), anthocyanins (8), capacity of electron flow (PSI) (6), carotenoids (42), chlorophyll a content (59), chlorophyll b content (54), effective quantum yield (4), glycolate (1), gross photosynthetic rate (7), initial slope of rapid light curve (5), intrinsic quantum yield (1), irradiance at the onset of saturation (Ek) (4), maximum fluorescence level (Fm) (3), maximum photochemical quantum yield (Fv/Fm) (30), maximum photosynthetic rate (4), minimum fluorescence level (F0) (1), nadh enzyme (1), neoxanthin (1), net photosynthesis (10), non-photochemical fluorescence quenching (npq) (9), phosphoenolpyruvate carboxylase (1), photosynthetic CO2 response curve (1), photosynthetic HCO3^-^ response curve (1), quantum yield y (19), relaxation (1), rubisco activity (5), saturating irradiance (2), total chlorophyll (54), violaxanthin (2), half saturation constant (Km) (1) |
|  | Respiration | oxygen consumption (11) |
|  | Water | osmotic potential (1) |
| Morphological | Biomass Distribution | allocation ratio (1), leaf mass fraction (2), leaf mass ratio (1), leaf mass per area (4), root mass fraction (2), root mass ratio (1), root % volume (1), specific leaf ratio (1), stem dry matter content (1), stem mass fraction (1), total mass ratio (1) |
|  | Competitive Ability | dominance ratio (1) |
|  | Count | branch number (22), bud number (1), fraction of leaves (1), leaf bundle count (5), leaf number (8), number of crossings (1), number of forks (1), number of root tips (1), ramet number (3), root number (5), rosette number (1), turion number (1), whorl number (1) |
|  | Growth Rate | growth inhibition rate (1), growth rate (69) |
|  | Mass | ash weight (2), dry weight (42), flower biomass (1), fresh weight (42), individual branch biomass (1), leaf mass (2), leaf weight (2), root dry weight (1), root fresh weight (1), root mass (1), root weight (8), seed biomass (1), stem mass (1), stolon weight (1), tuber depletion (ash-free dw) (1), tuber weight (1) |
|  | Phenology | flowering date (1) |
|  | Reproductive Success | autofragmentation (1), flowering branch number (2), germination rate (1), seed setting rate (1) |
|  | Resistance | permeability (1), sclerophylly index (1) |
|  | Size | branch length (3), internode length (4), leaf length (7), leaf thickness (1), leaf width (3), plant height (1), rhizoid length (1), root diameter (1), root length (20), root surface area (1), seedling height (1), specific leaf area (sla) (5), stem length (1), total projected area (1), length (3) |
|  | Tissue density | dry matter content (dmc) (3), leaf dry matter content (ldmc) (5), root dry matter content (rdmc) (3), root moisture content (rmc) (1) |
|  | Visual damage | appearance (1), cell structure (8), chloroplast structure (2), dead leaf tissue (3), leaf structure (1), mitochondria structure (1) |

Table S3: Summary table of the modelled physiological end point changes with single stress. ID_XP: Experiment ID, SP_ID: Species ID, ID_XP:StressorUnit : Stressor identity nested within the experiment (as applied stressor levels and origin were different). *σ*^2^: term variance I2: heterogeneity index.

| **Physiological end point** | **k** | **estimate** | **ci.lb** | **ci.ub** | **pval** | **Percent change** | **Model expression** | *σ*^2^**_ID_XP** | *σ*^2^**_SP_ID** | *σ*^2^**_nested** | $I^{2}$**_total** | $I^{2}$**_ID_XP** | $I^{2}$**_SP_ID** | $I^{2}$**_nested** |
| --- | --- | --- | --- | --- | --- | --- | --- | --- | --- | --- | --- | --- | --- | --- |
| **H2O2** | 27 | 0.418 | 0.126 | 0.709 | 0.005 | 51.834 | ID_XP | 0.247 | <10^-6^ | <10^-6^ | 0.551 | 0.551 | <10^-6^ | <10^-6^ |
| **CAT** | 70 | 0.403 | 0.175 | 0.631 | 0.001 | 49.579 | ID_XP | 0.370 | <10^-6^ | <10^-6^ | 0.964 | 0.964 | <10^-6^ | <10^-6^ |
| **POD** | 84 | 0.309 | 0.072 | 0.547 | 0.011 | 36.253 | ID_XP+ID_XP:StressorUnit | 0.213 | <10^-6^ | 0.213 | 0.924 | 0.462 | <10^-6^ | <10^-6^ |
| **SOD** | 99 | 0.303 | 0.184 | 0.423 | <10^-4^ | 35.445 | ID_XP+SP_ID+ID_XP:StressorUnit | 0.023 | 0.001 | 0.023 | 0.682 | 0.332 | 0.019 | <10^-6^ |
| **MDA** | 114 | 0.218 | 0.129 | 0.308 | <10^-4^ | 24.414 | ID_XP | 0.089 | <10^-6^ | <10^-6^ | 0.847 | 0.847 | <10^-6^ | <10^-6^ |
| **P** | 42 | 0.184 | 0.002 | 0.366 | 0.048 | 20.181 | ID_XP+ID_XP:StressorUnit | 0.047 | <10^-6^ | 0.047 | 0.834 | 0.417 | <10^-6^ | <10^-6^ |
| **N** | 92 | 0.109 | -0.014 | 0.233 | 0.083 | 11.532 | ID_XP+SP_ID+ID_XP:StressorUnit | 0.020 | 0.011 | 0.02 | 0.812 | 0.317 | 0.177 | <10^-6^ |
| **C** | 55 | -0.010 | -0.057 | 0.038 | 0.689 | -0.968 | ID_XP+ID_XP:StressorUnit | 0.005 | <10^-6^ | 0.005 | 0.754 | 0.377 | <10^-6^ | <10^-6^ |
| **Fv/Fm** | 74 | -0.059 | -0.121 | 0.002 | 0.058 | -5.764 | SP_ID | 0 | 0.013 | <10^-6^ | 0.605 | <10^-6^ | 0.605 | <10^-6^ |
| **Chl** | 113 | -0.096 | -0.259 | 0.068 | 0.251 | -9.131 | ID_XP+SP_ID+ID_XP:StressorUnit | 0.009 | 0.039 | 0.009 | 0.405 | 0.064 | 0.277 | <10^-6^ |
| **Soluble protein** | 46 | -0.151 | -0.294 | -0.008 | 0.038 | -14.046 | ID_XP+ID_XP:StressorUnit | 0.006 | <10^-6^ | 0.006 | 0.352 | 0.176 | <10^-6^ | <10^-6^ |
| **Chl b** | 116 | -0.228 | -0.365 | -0.090 | <10^-4^ | -20.379 | ID_XP+ID_XP:StressorUnit | 0.063 | <10^-6^ | 0.063 | 0.672 | 0.336 | <10^-6^ | <10^-6^ |
| **Chl a** | 124 | -0.285 | -0.436 | -0.134 | <10^-4^ | -24.821 | ID_XP+ID_XP:StressorUnit | 0.073 | <10^-6^ | 0.073 | 0.852 | 0.426 | <10^-6^ | <10^-6^ |

Table S4: Summary table of the modelled physiological response of submerged macrophytes to single stress (k: number of effect sizes, n: number of studies, ci.lb – ci.ub: lower and upper bound of the confidence interval, *σ*^2^: term variance, $I^{2}:$ heterogeneity index.

| **Stressor class** | **k** | **n** | **estimate** | **ci.lb** | **ci.ub** | **pval** | **Percent change** | **Model expression** | *σ*^2^**_ID_XP** | *σ*^2^**_SP_ID** | *σ*^2^**_Physio_ID** | *σ*^2^**_nested** | $I^{2}$**_total** | $I^{2}$**_ID_XP** | $I^{2}$**_SP_ID** | $I^{2}$**_Physio_ID** | $I^{2}$**_nested** |
| --- | --- | --- | --- | --- | --- | --- | --- | --- | --- | --- | --- | --- | --- | --- | --- | --- | --- |
| **Shading** | 93 | 31 | 0.15 | -0.06 | 0.35 | 0.17 | 15.62 | ID_XP+SP_ID+Physio_ID+ID_XP:StressorUnit | 0.048 | 0.064 | 0.039 | 0.048 | 0.909 | 0.219 | 0.291 | 0.180 | 0.219 |
| **Nutrient enrichment** | 200 | 52 | 0.07 | -0.04 | 0.18 | 0.21 | 7.18 | ID_XP | 0.152 | <10^-6^ | <10^-6^ | <10^-6^ | 0.788 | 0.788 | <10^-6^ | <10^-6^ | <10^-6^ |
| **Carbon dioxide** | 42 | 11 | -0.08 | -0.24 | 0.09 | 0.37 | -7.29 | ID_XP+SP_ID+Physio_ID+ID_XP:StressorUnit | 0.017 | <10^-6^ | 0.012 | 0.017 | 0.773 | 0.286 | 0.002 | 0.198 | 0.286 |
| **Microplastics** | 85 | 10 | -0.13 | -0.22 | -0.03 | 0.01 | -11.77 | ID_XP | 0.021 | <10^-6^ | <10^-6^ | <10^-6^ | 0.562 | 0.562 | <10^-6^ | <10^-6^ | <10^-6^ |
| **Cyanotoxin** | 33 | 7 | -0.22 | -0.48 | 0.04 | 0.09 | -20.02 | ID_XP+SP_ID+Physio_ID | 0.013 | 0.029 | 0.060 | <10^-6^ | 0.891 | 0.115 | 0.256 | 0.520 | <10^-6^ |
| **Warming** | 61 | 21 | -0.22 | -0.59 | 0.15 | 0.23 | -20.11 | ID_XP+SP_ID+Physio_ID+ID_XP:StressorUnit | 0.068 | 0.108 | 0.163 | 0.068 | 0.830 | 0.139 | 0.220 | 0.331 | 0.139 |
| **Antibiotic** | 40 | 7 | -0.24 | -0.96 | 0.49 | 0.52 | -20.96 | ID_XP+SP_ID+Physio_ID+ID_XP:StressorUnit | 0.246 | 0.049 | 0.379 | 0.246 | 0.981 | 0.262 | 0.053 | 0.404 | 0.262 |
| **Nanoparticles** | 30 | 6 | -0.25 | -0.47 | -0.02 | 0.03 | -21.75 | Physio_ID | <10^-6^ | <10^-6^ | 0.118 | <10^-6^ | 0.873 | <10^-6^ | <10^-6^ | 0.873 | <10^-6^ |
| **Other chemicals** | 62 | 7 | -0.29 | -0.65 | 0.07 | 0.11 | -25.24 | ID_XP+SP_ID+Physio_ID+ID_XP:StressorUnit | 0.029 | 0.001 | 0.202 | 0.028 | 0.960 | 0.108 | 0.003 | 0.745 | 0.103 |
| **PFASs** | 44 | 8 | -0.35 | -0.58 | -0.12 | 0.00 | -29.40 | Physio_ID | <10^-6^ | <10^-6^ | 0.106 | <10^-6^ | 0.834 | <10^-6^ | <10^-6^ | 0.834 | <10^-6^ |
| **Metals** | 221 | 34 | -0.35 | -0.53 | -0.17 | 0.00 | -29.51 | ID_XP+SP_ID+Physio_ID+ID_XP:StressorUnit | 0.002 | 0.034 | 0.030 | 0.002 | 0.614 | 0.022 | 0.304 | 0.266 | 0.022 |

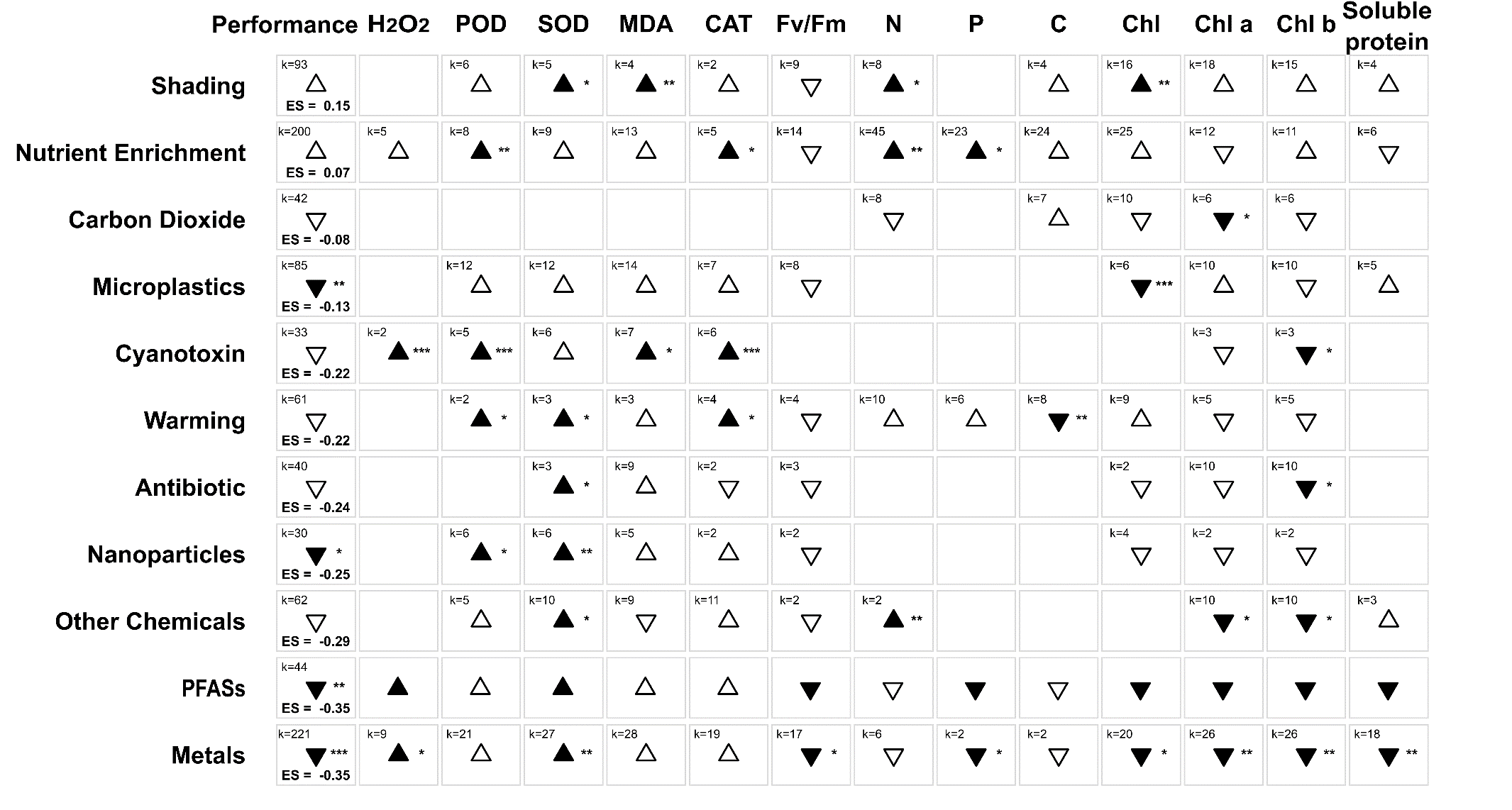

Fig. S6: Physiological responses of submerged macrophytes to main single stress classes. The overall performance is the modelled estimate across physiological end points. ES is the estimated effect size and k represents the number of effect sizes. Triangles demonstrate a change of estimate (up is more, down is less). Dark triangles demonstrate a significant change and hollow triangles a non-significant change (p value > 0.05). significance stars (*** = p < 0.001; **= p < 0.01; * = p < 0.05).

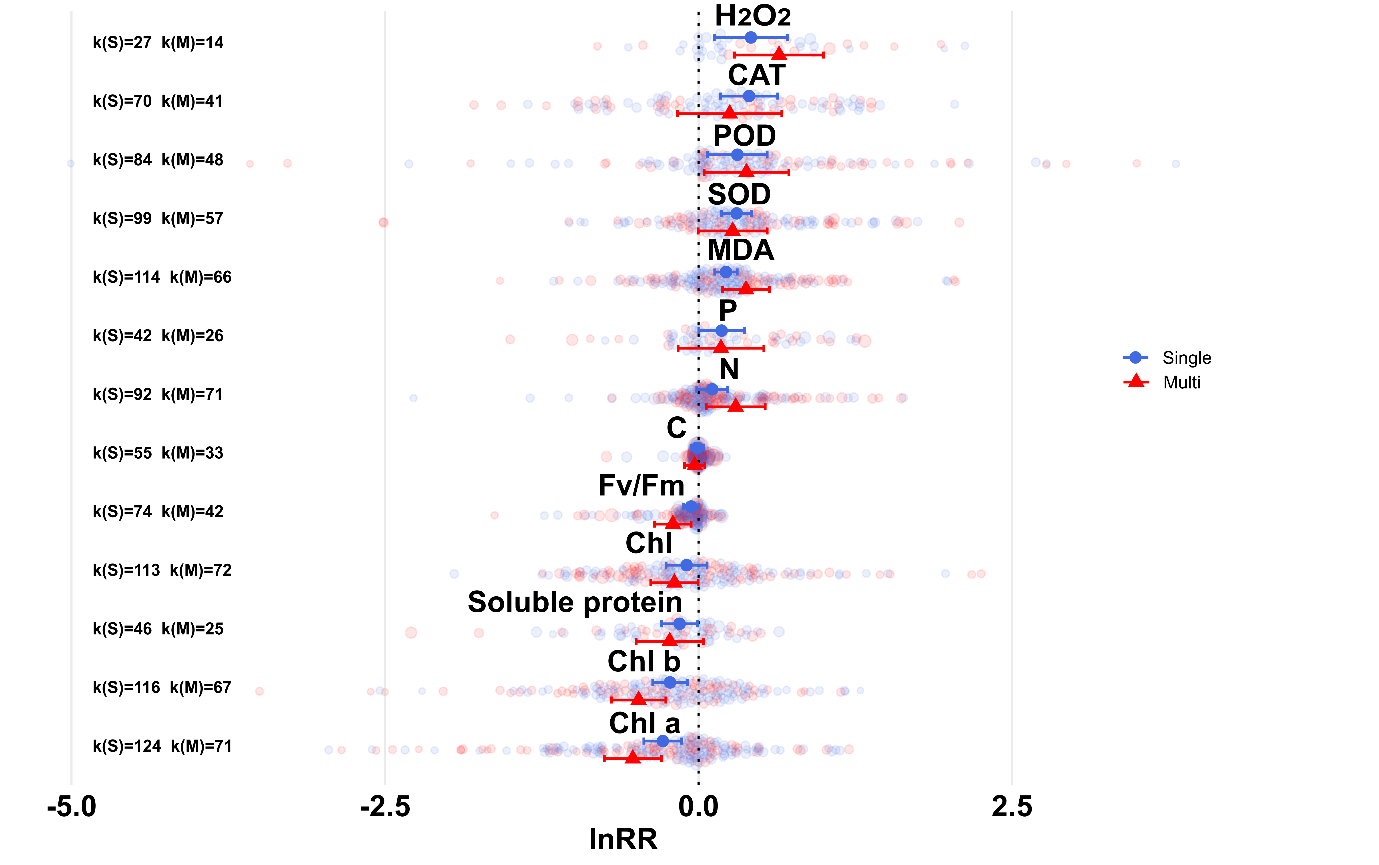

Fig. S7: Orchard plot showing lnRR estimates of the most frequently measured physiological endpoints changes under single (S, blue) and multiple stressors (M, red). Horizontal lines indicate 95% confidence intervals, and individual lnRR effect sizes are shown as points coloured by stressor combi0tion. Endpoint-specific differences between single and multiple stressor effects were assessed using Wald tests on pooled estimates with no significance obtained.

Table S5: Comparison of multi- and single-stressor effect sizes (lnRR) across endpoint classes using Welch’s unequal-variances t-tests.

| **Endpoint** | **Multi (lnRR ± SE)** | **Single (lnRR ± SE)** | **t** | **df** | **p-value** |
| --- | --- | --- | --- | --- | --- |
| H₂O₂ | 0.642 ± 0.181 | 0.418 ± 0.149 | -0.96 | 29.74 | 0.346 |
| N | 0.296 ± 0.120 | 0.109 ± 0.063 | -1.38 | 107.73 | 0.171 |
| MDA | 0.378 ± 0.096 | 0.218 ± 0.046 | -1.51 | 95.02 | 0.135 |
| POD | 0.382 ± 0.172 | 0.309 ± 0.121 | -0.34 | 92.1 | 0.731 |
| P | 0.179 ± 0.174 | 0.184 ± 0.093 | 0.03 | 39.23 | 0.979 |
| C | -0.030 ± 0.041 | -0.010 ± 0.024 | 0.43 | 54.41 | 0.666 |
| SOD | 0.272 ± 0.140 | 0.303 ± 0.061 | 0.21 | 77.62 | 0.837 |
| Soluble protein | -0.230 ± 0.137 | -0.151 ± 0.073 | 0.51 | 38 | 0.615 |
| Chl | -0.192 ± 0.096 | -0.096 ± 0.083 | 0.76 | 160.49 | 0.449 |
| F_v_/F_m_ | -0.205 ± 0.074 | -0.059 ± 0.031 | 1.81 | 55.93 | 0.076 |
| CAT | 0.248 ± 0.212 | 0.403 ± 0.116 | 0.64 | 64.37 | 0.526 |
| Chl a | -0.524 ± 0.116 | -0.285 ± 0.077 | 1.71 | 130.85 | 0.089 |
| Chl b | -0.478 ± 0.111 | -0.228 ± 0.070 | 1.91 | 118.7 | 0.059 |

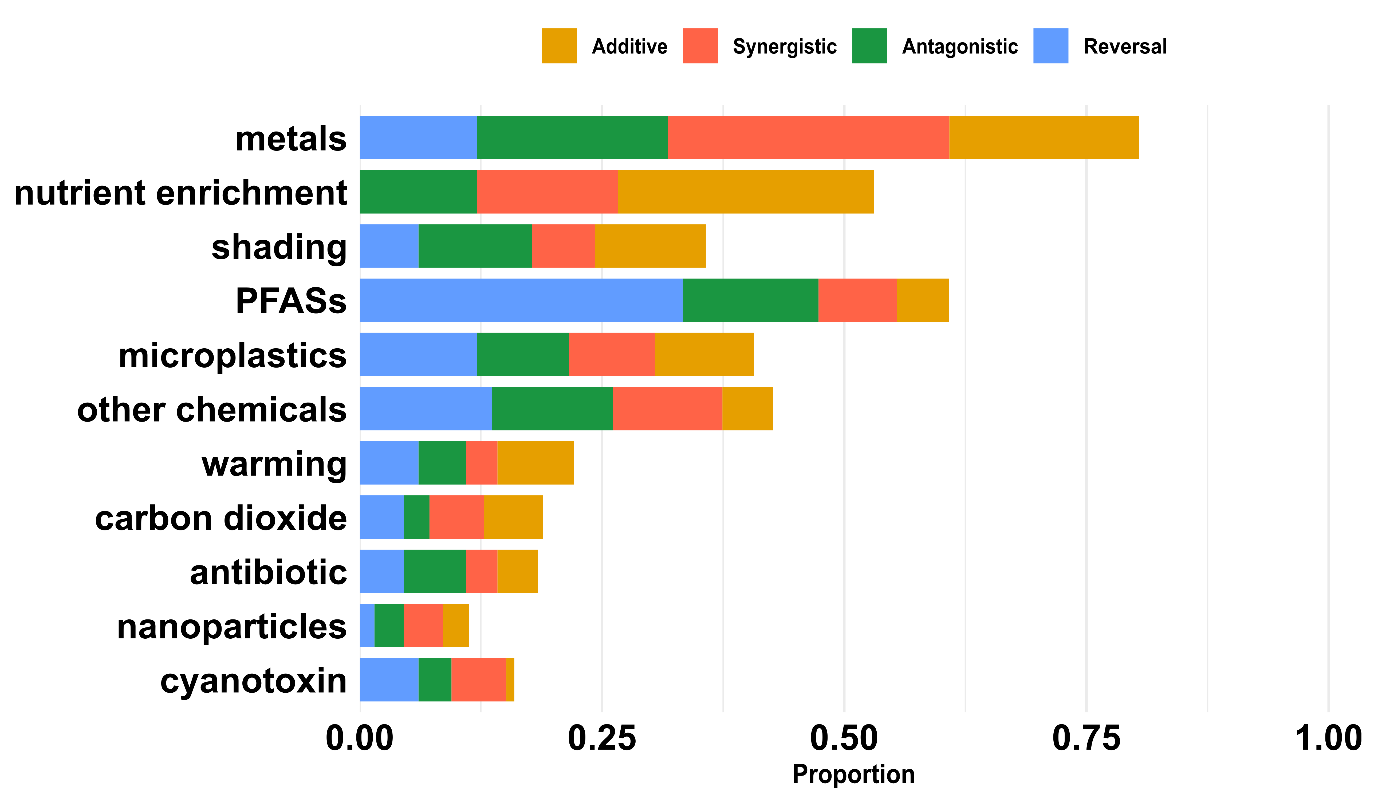

Fig. S8: Distribution of interaction types across major stressors as a function of combination proportions, illustrating the contribution of individual stressors to each interaction class.

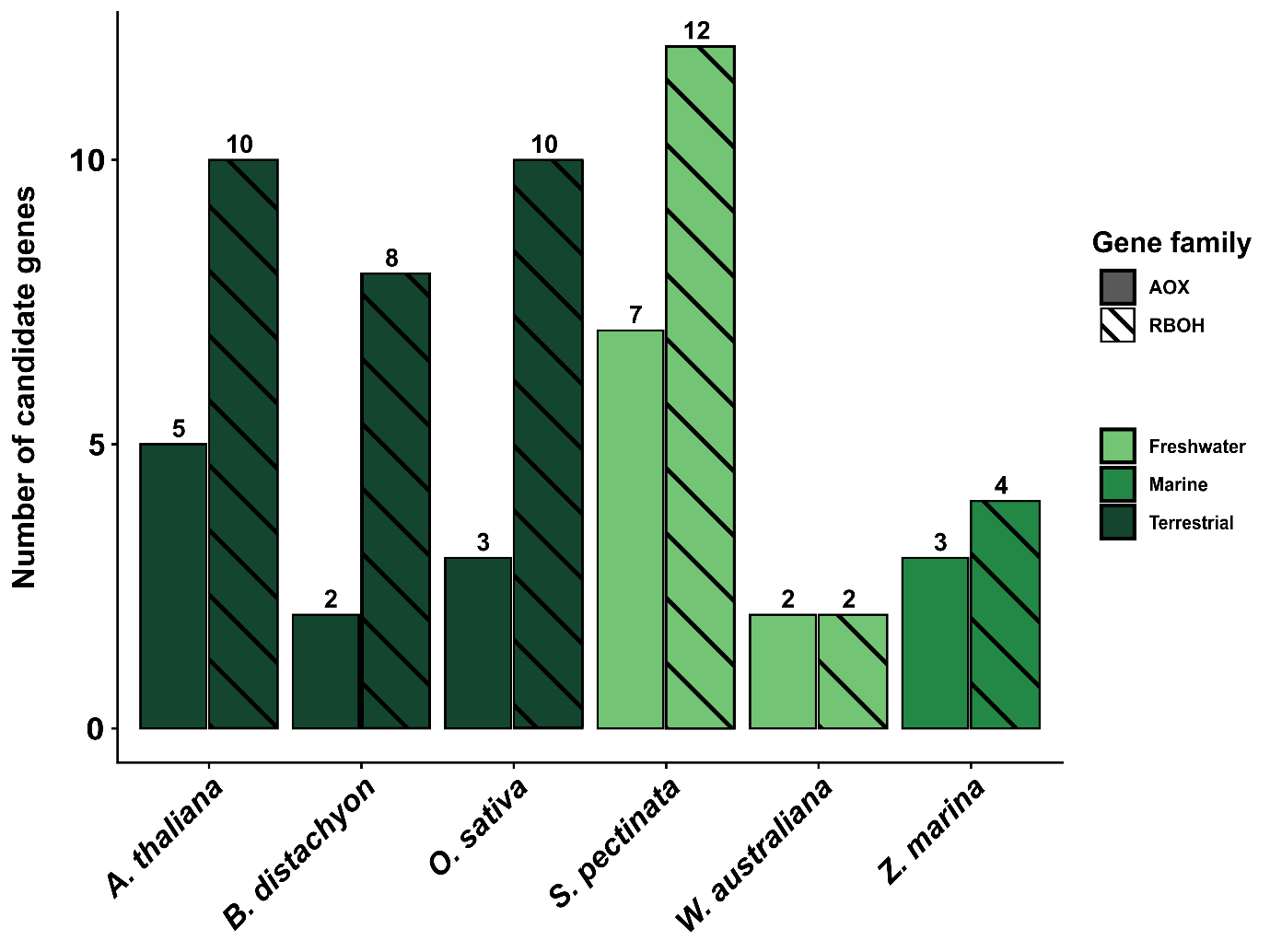

Fig. S9: Comparative abundance of AOX and RBOH gene families across selected terrestrial, marine, and freshwater angiosperms.

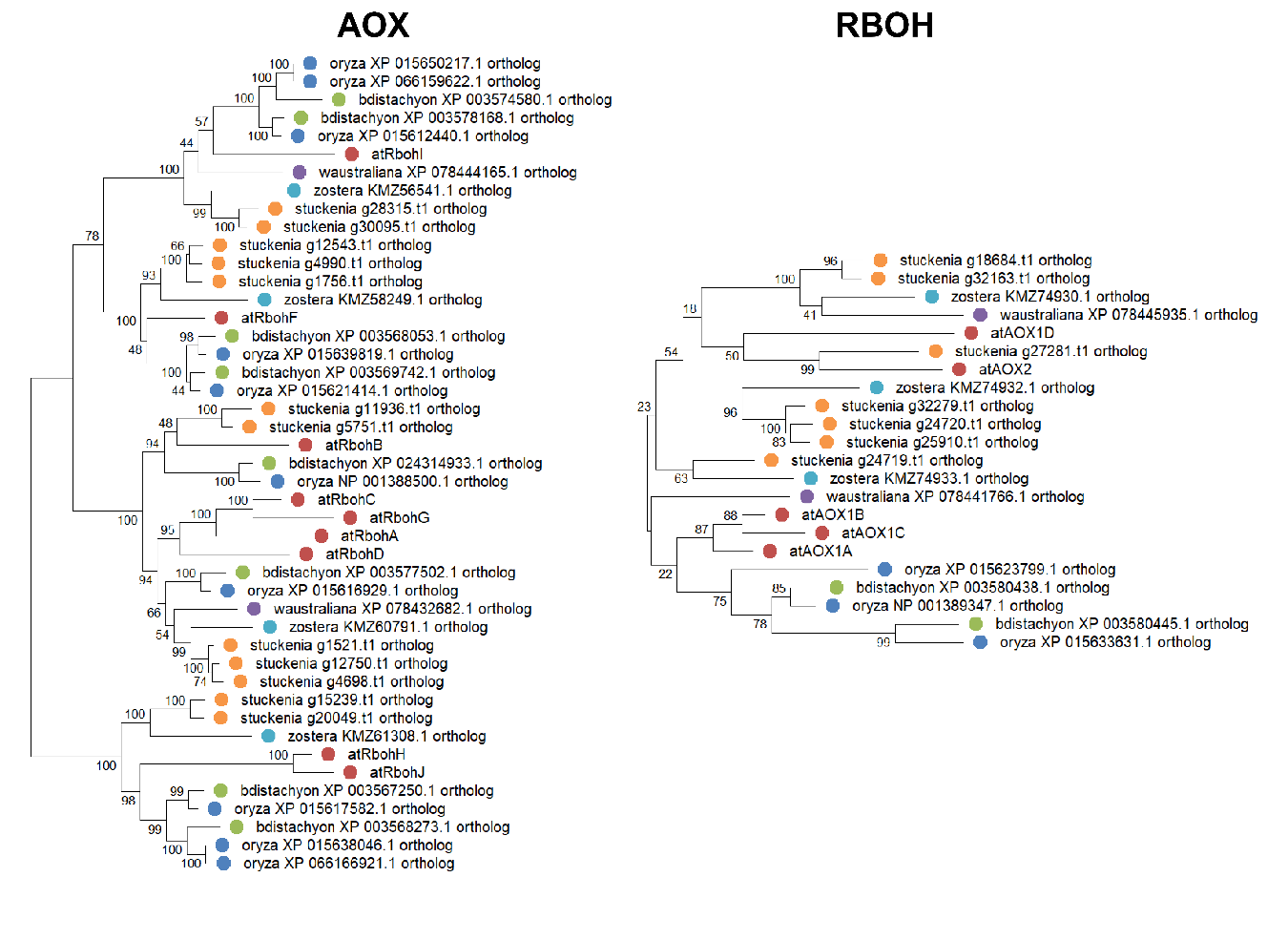
Fig. S10: Phylogenetic analysis of candidate gene families: maximum-likelihood tree of putative AOX and RBOH protein sequences.

Table S6: R packages used for the information synthesis, meta-analysis and figure production.

| **Package** | **Version** | **Citation (Author, Year)** |
| --- | --- | --- |
| dplyr | 1.2.0 | Wickham et al., 2026 |
| extrafont | 0.2 | Chang, 2025 |
| ggbeeswarm | 0.7.3 | Clarke et al., 2025 |
| ggplot2 | 4.0.2 | Wickham, 2016 |
| ggraph | 2.2.2 | Pedersen, 2025 |
| ggrepel | 0.9.6 | Slowikowski, 2024 |
| ggtext | 0.1.2 | Wilke & Wiernik, 2022 |
| igraph | 2.2.2 | Csárdi & Nepusz, 2006 |
| Matrix | 1.7.4 | Bates et al., 2025 |
| metafor | 4.8.0 | Viechtbauer, 2010 |
| patchwork | 1.3.2 | Pedersen, 2025 |
| purrr | 1.2.1 | Wickham & Henry, 2026 |
| raster | 3.6.32 | Hijmans, 2025 |
| RColorBrewer | 1.1.3 | Neuwirth, 2022 |
| readr | 2.1.6 | Wickham et al., 2025 |
| readxl | 1.4.5 | Wickham & Bryan, 2025 |
| reshape2 | 1.4.5 | Wickham, 2007 |
| scales | 1.4.0 | Wickham et al., 2025 |
| sf | 1.0.24 | Pebesma, 2018 |
| spData | 2.3.4 | Bivand et al., 2025 |
| stringr | 1.6.0 | Wickham, 2025 |
| svglite | 2.2.2 | Wickham et al., 2025 |
| taxize | 0.10.1 | Chamberlain & Szöcs, 2013 |
| tidyr | 1.3.2 | Wickham et al., 2025 |
| tidyverse | 2.0.0 | Wickham et al., 2019 |
| viridis | 0.6.5 | Garnier et al., 2024 |
